# Improved Protein Semi-Synthesis Enables Biophysical Studies of Thioamide Destabilization of β-Sheet Interactions

**DOI:** 10.64898/2026.07.30.741859

**Authors:** Evan S. K. Yanagawa, Kristen E. Fiore, Denver Y. Francis, Aiden Lesneski, Yanan Chang, Benjamin Roose, David Christianson, Kohei Sato, E. James Petersson

**Affiliations:** Department of Chemistry, School of Arts and Sciences, University of Pennsylvania; 231 S. 34th Street, Philadelphia, PA 19104, USA; Department of Engineering, Graduate School of Integrated Science and Technology, Shizuoka University, 3-5-1 Johoku, Hamamatsu, Shizuoka 432-8561, Japan; Department of Biochemistry and Biophysics, Perelman School of Medicine, University of Pennsylvania, 421 Curie Boulevard, Philadelphia, PA 19104, USA

## Abstract

Thioamides are natural post-translational modifications of the peptide backbone and can be introduced synthetically to probe protein folding or functionalize peptides for translational applications. In this work, we demonstrate that thioamide-containing peptides with C-terminal thioesters can be efficiently generated using Knorr pyrazole activation and used in subsequent native chemical ligation reactions to generate thioamide containing proteins. We compare this method to acyl azide activation and find that both routes provide similar yields. We also investigate ultrasound-mediated desulfurization of the ligation site cysteine for potential advantages over chemical radical initiators. Scaling up our syntheses allows us to study thioamide perturbations to the β-sheet region of the B1 domain of protein G (GB1) as well as β-strand interactions in amyloid fibrils of the Parkinson’s disease protein α-synuclein. In both contexts, we observe dramatic destabilization of the β-sheet networks, manifested in decreased GB1 thermal stability and altered folding and slowed aggregation of α-synuclein. These findings illustrate the impact that a single atom substitution can have on cooperative hydrogen bonding networks and prompt future study of both systems.

## Introduction

Thioamides are conservative backbone modifications in which the carbonyl oxygen of an amide bond is replaced by sulfur, producing a thiocarbonyl (C=S).^1–8^ This single-atom substitution introduces distinct physicochemical changes that can be highly informative for biophysical studies.^2^ Compared with “oxoamides,” thioamides possess a longer and more polarizable C=X bond, increased rotational rigidity about the C–N bond, and altered electronic structure, which can influence local conformational preferences and backbone dynamics.^2, 9–11^ In addition, thioamide N–H groups are more acidic and function as stronger hydrogen-bond donors, whereas the sulfur atom is typically a weaker hydrogen-bond acceptor than oxygen, enabling selective perturbation of hydrogen-bond networks that stabilize protein secondary structure.^12, 13^ These effects can be applied to increase resistance to proteolytic degradation or modulate peptide stability and bioactivity, reflecting the altered electronic structure and reduced susceptibility of the C=S bond to enzymatic hydrolysis.^1, 5, 7, 14–16^ Nature also harnesses the thioamide, incorporating it into a few classes of peptide natural products and into some *bona fide* proteins, two of which have been structurally characterized.^2, 17, 18^ The roles of the thioamides in these proteins are not fully understood, further motivating the study of thioamides in human-made systems.

The electronic structure differences between thioamides and oxoamides provide characteristic spectroscopic signatures, including red-shifted UV absorption and distinctive NMR and IR features, as well as the ability to quench fluorophores through Förster resonance energy transfer (FRET) or photo-induced electron transfer (PET) mechanisms.^19–25^ These unique properties position thioamides to serve as minimally perturbing probes of local structure and environment, which makes them valuable tools in chemical biology and protein biophysics. Our laboratory has long been interested in harnessing these properties for the study of conformational changes in complex systems of biomedical interest, such as the disordered and amyloidogenic protein α-synuclein (αS), which is associated with Parkinson’s disease.^26–28^ We have also sought to establish benchmarks for the impact of thioamides on protein folding through the study of model protein systems with well-characterized structures and folding transitions.^29, 30^ Both types of investigations often involve large quantities of thioamide proteins, motivating the exploration of alternative native chemical ligation (NCL) approaches to improve yields.

In the case of αS aggregation, studies are typically performed at 100 μM concentrations in mL-scale volumes, requiring quantities of thioamide αS that have been difficult to access. To monitor conformational changes in αS, we incorporated thioamides to serve as minimally-perturbing FRET or PET quenching probes. For example, we utilized the ability of the thioamide to quench Trp fluorescence through PET to monitor early misfolding events in αS aggregation.^26^ Since αS contains no native Cys residues, the thiovaline 3 variant (V^S^_3_, thioamide denoted as superscript “S”) was synthesized as a S_9_C mutant by NCL of a αS_1-8_-V^S^_3_ thioester with αS_9-140_-S_9_C. The resulting thioamide construct was then mixed 1:10 with wild type (WT) αS for aggregation so that fluorescence changes would arise only from intramolecular conformational changes upon formation of oligomers or fibrils. Subsequent studies used unnatural amino acid mutagenesis or ligation to incorporate other fluorescent donors to pair with the thioamide as FRET or PET partners.^27, 31, 32^ The use of 10% mixtures of thioamide αS with WT αS allowed us to isolate intramolecular conformational changes, but was also a strategy of convenience, since quantities of thioamide αS were limited. Improvements in thioamide NCL would enable broader use of thioamides in these aggregation experiments.

In parallel with investigations of αS, we conducted thioamide benchmarking studies using a series of small proteins representing canonical secondary structures, including the compact immunoglobulin-binding B1 domain of protein G from *Streptococcus* bacterium (GB1). The 56 amino acid GB1 protein adopts a tight fold comprising a four-stranded β-sheet packed against a protein-spanning α-helix and has been used for a variety of biophysical and protein folding studies, including substitutions of the peptide backbone.^33–36^ We used NCL to generate three single-thioamide GB1 variants positioning the modification at L_5_, I_6_, or L_7_—sites on the same β-strand within the core of GB1.^30^ Thermodynamic stability measurements of WT GB1 and the three thioamide variants were made using CD melt experiments. Although positioned in similar environments, L^S^_5_, I^S^_6_, and L^S^_7_ imparted markedly different effects on protein stability differing by 2 kcal/mol. We attempted to rationalize these effects using available high-resolution structural data on WT GB1 (Protein Data Bank, or PDB, ID 2QMT),^37^ but we lacked structural data on the thioamide variants. In contrast, when we studied a model β-hairpin peptide, we were able to produce multi-mg quantities of the thioamide constructs through peptide synthesis, enabling analysis by NMR as well as CD.^29^ For the β-hairpin peptide, we found that thioamide stabilization as a hydrogen bond donor or destabilization as a hydrogen bond acceptor was position dependent, and NMR-based structural models helped to explain these subtle effects. This study highlighted the value of being able to produce sufficient quantities of protein for structural characterization and prompted us to seek improved methods for the synthesis of thioamide variants of proteins like GB1 and αS. Moreover, as we gained greater understanding of the effects of thioamides in β-sheet model systems, we were increasingly interested in studying their effects on the αS fibril core, consisting of repeated β-sheet elements. Such experiments would require the thioamide to be present in 100% of the monomers in fibrils, again necessitating higher yielding NCL methods.

The yields of our syntheses of thioamide proteins have primarily depended on the yields of the thioamide-containing peptides obtained through solid phase peptide synthesis. In the investigations of GB1 and αS noted above, these required the synthesis of thioamidated N-terminal protein fragments activated with a C-terminal thioester for NCL. We have previously explored a number of strategies for the synthesis of such fragments. Initial attempts used PyBOP activation. However, this method is hampered by epimerization at the α-carbon of the C-terminal amino acid and poor solubility of the sidechain-protected peptides.^26, 27, 31^ Utilizing a C-terminal cysteinyl prolyl ester with unprotected peptides avoided both of these concerns.^38^ However, this method is inefficient since it entails synthesis of a dipeptide with the desired C-terminal amino acid, inefficient coupling of the sterically hindered Pro to glycolic acid, and overall instability at the ester. We have also explored use of the Dawson 3,4-diaminobenzoyl (Dbz) resin.^39^ This chemistry was confirmed to be compatible with a thioamide-containing peptide, but a Pd-deprotection step sometimes used in Dbz activation degraded the thioamide.^40^ More recently, we have focused on methods using C-terminal acyl hydrazides.^30, 41, 42^

Inspired by the need for NCL strategies that yield sufficient quantities of thioamide proteins for biophysical studies, we wished to explore various C-terminal acyl hydrazide activation strategies to generate reactive thioesters *in situ*.^41, 42^ In NCL methods popularized by Liu, acyl hydrazides are activated to form C-terminal thioesters for NCL by the addition of oxidants.^43^ A typical oxidant is sodium nitrite, which under acidic conditions will convert a C-terminal acyl hydrazide to an acyl azide. This species is activated for exchange with thiols to form the C-terminal thioester for NCL (**Figure 1**), an approach that we have applied to thioamide protein synthesis.^30, 41, 42^ However, we have struggled to achieve isolated yields greater than 30%. Potential sources of side-products include the formation of thioamide *S*-nitroso species with subsequent degradation.^44^ Thus, we were intrigued to try activation with other non-redox active reagents. We have therefore investigated activation to the acyl Knorr pyrazole with acetyl acetone as demonstrated by Dawson and coworkers (**Figure 1**). However, we had some concerns that the nucleophilicity of the thioamide sulfur could result in off-pathway reactions with the diketone reagent resulting in S-to-O exchange or peptide hydrolysis at the thioamide. Thus, we tested Knorr pyrazole formation with a familiar N-terminal GB1 fragment.

**Figure 1.**
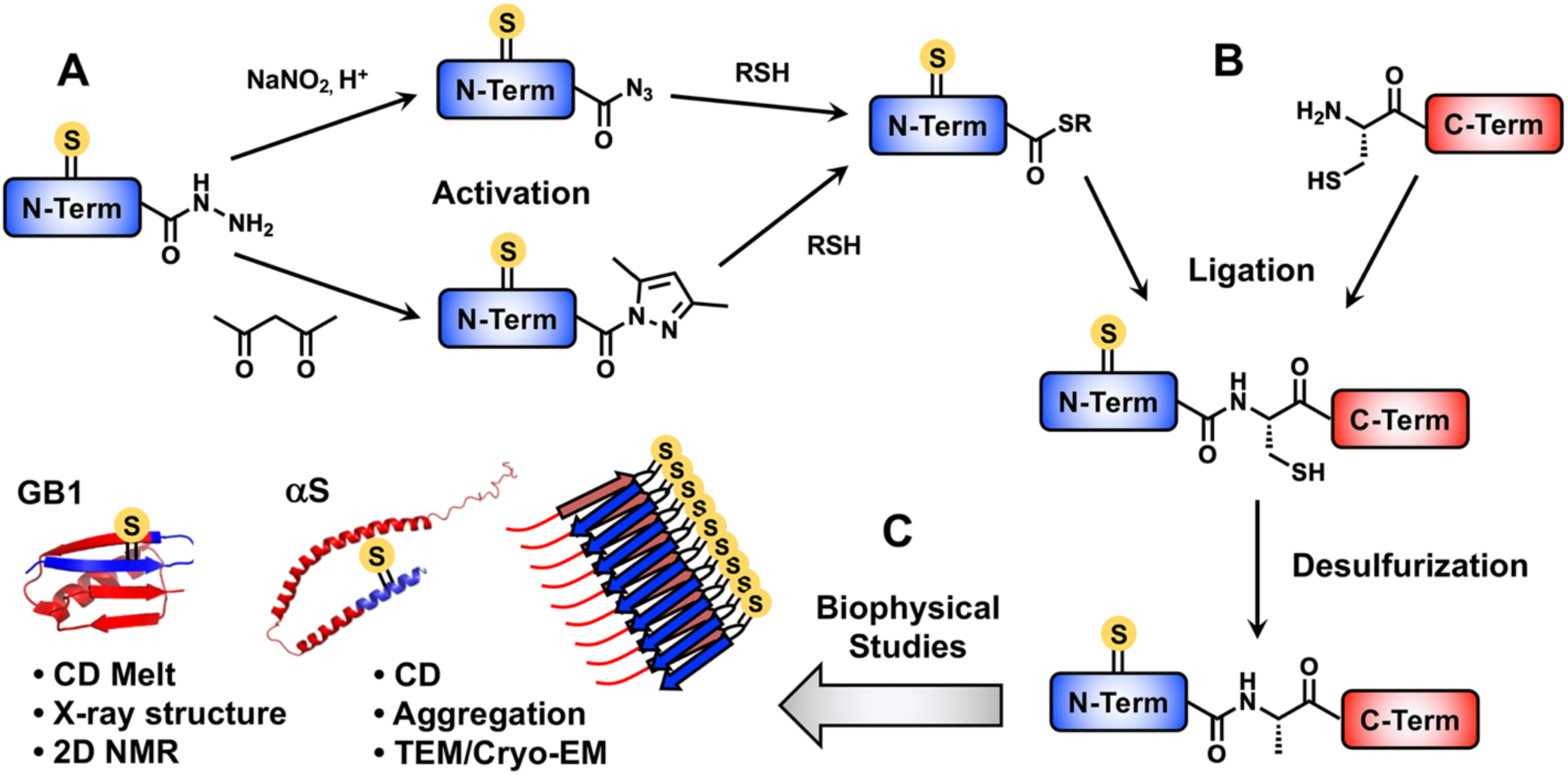
Methods for synthesis of thioamide proteins and biophysical analysis of model proteins GB1 and αS. **A**) Synthesis of N-terminal protein fragments (N-Term) with C-terminal thioesters from acyl hydrazides via acyl azides or Knorr pyrazoles (this work). **B**) Ligation of the N-term to an expressed C-terminal fragment (C-Term) with a cysteine, followed by desulfurization using radical initiators or ultrasound (this work), provides the full-length thioamide protein. **C**) Thioamide proteins can be analyzed by biophysical and structural methods for comparison to the oxoamide parent protein. GB1 CD, X-ray, and NMR, and αS aggregation studies are attempted in this work.

Here we report direct comparison of Knorr pyrazole and acyl azide thioester formation approaches in the semi-synthesis of GB1 and show that we are able to produce multi-mg quantities of protein, enabling NMR studies to investigate a strongly perturbing thioamide position. We also apply both approaches to thioamide αS synthesis, allowing us to perform aggregation experiments with up to 100% modified protein. Surprisingly, we observe a dramatic slowing of aggregation for a thioamide at an N-terminal position not expected to form part of the amyloid fibril core. These studies underscore the impact that thioamide substitution can have in β-sheet networks in spite of its many shared features with oxoamide peptide bonds.

## Materials and Methods

General information for reagents and instruments, synthetic procedures for thioamide precursors, peptide synthesis and purification, protein expression and purification are given in Supporting Information.

### Acyl Azide Activation and Ligation

The activation and NCL buffers were made the day before and argon purged prior to starting the ligation (activation buffer: 6 M guanidine (Gdn) HCl, 0.2 M NaH_2_PO_4_ pH 3.0, NCL buffer: 6 M Gdn HCl, 0.2 M phosphate, 0.2 M 4-mercaptophenylacetic acid (MPAA) pH 7.0). The acyl hydrazide peptide was dissolved in the activation buffer to achieve a 2 mM peptide concentration. The peptide was left stirring in a -15 °C in an NaCl/ice bath in a Dewar bowl. After the peptide was cooled, 10 equivalents of sodium nitrate (from a 1 M stock in sterile Milli-Q water) was added and the peptide was left to stir for 15 minutes. In the meanwhile, the N-terminal cysteine containing protein was dissolved in NCL buffer to achieve a ratio of 16:1 MPAA/ NaNO_2_. After the 15-minute incubation, the protein was added to the chilled peptide. The pH was monitored and carefully adjusted to ∼ pH 7.0. The reaction was monitored by MALDI MS and analytical RP-HPLC. A freshly prepared solution of TCEP neutral was added to the reaction after 3-4 hours to a final concentration of 50 mM TCEP. After the reaction was determined complete by MALDI MS, the NCL was slowly diluted with Milli-Q water and additional TCEP was added before dialysis and/or purification. Additional details are provided in Supporting Information.

### Knorr Pyrazole Activation and Ligation

The acyl hydrazide peptide was dissolved in 6 M Gdn HCl to 2 mM concentration. Solid MPAA was weighed out to achieve a 0.2 M MPAA solution and was carefully transferred to the peptide solution. From a 100 mM stock in Milli-Q water, acyl acetone (2.5 eq, 0.975 g/mL) was added to the peptide and allowed to stir at room temperature. The reaction was monitored by MALDI-TOF MS via a 100x dilution in Milli-Q water with 0.1 % TFA. After 3 hours, the thioester formation was determined complete by analytical. The NCL buffer (6 M Gdn HCl, 0.2 M Na_2_HPO_4_ pH 8.5) was purged with argon. The N-terminal cysteine containing protein was dissolved in the NCL buffer to a concentration of 2 mM protein with a final concentration of 50 mM TCEP (from a 0.5 M TCEP neutral stock). The protein was added to the peptide and the pH was monitored and adjusted to pH 7.0-7.5. The ligation was allowed to proceed at room temperature overnight. After the reaction was determined complete by MALDI MS, the NCL was slowly diluted with Milli-Q water and additional TCEP was added before dialysis and/or purification. Additional details are provided in Supporting Information.

### GB1 Circular Dichroism (CD) Measurements

A dried aliquot of GB1 protein was denatured and refolded by dialysis as described in Supporting Information. CD experiments were performed on a Jasco J-1500 CD spectrometer with a 1 mm path length Helma 110-QS CD cuvette. Wavelength absorbance scans were performed at 25°C, scanning from 350 to 190 nm with a continuous scanning rate of 50 nm/min (bandwidth = 1 nm and data pitch = 1 nm). Thermal denaturation was monitored by changes in θ (mDeg) at 220 nm from 4-6 °C to 95 °C at a rate of 0.2 °C/min with an equilibration time of 10 seconds (interval = 1 °C, DIT = 8 sec and bandwidth = 1 nm). Additional experimental details, descriptions of data fitting, and primary data are included in Supporting Information.

### GB1 NMR Experiments

Lyophilized protein was dissolved in 50:50 v/v Milli-Q water/ acetonitrile and quantified by UV absorbance (ε_280nm_ = 9,970 M^-1^ cm^-1^ for GB1-K_13_A or ε_274nm_ = 10,169 M^-1^cm^-1^ for GB1-L^S^_5_K_13_A). An aliquot of 0.45-0.46 µmol was made and lyophilized. NMR samples were prepared by dissolving the lyophilized protein (≥ 95% purity) in 0.5-0.6 mL of 50 mM NaH_2_PO_4_ pH 5.3 (9:1 v/v Milli-Q H_2_O/ D_2_O) to 0.8 mM concentration. 2-Dimethyl-2-silapentane-5-sulfonate (DSS) was added to each NMR sample as the internal reference (from a 1 mg/mL stock in sterile Milli-Q water to ∼30 μM final concentration). NMR data were collected on a Bruker AVANCE NEO 600 MHz spectrometer. Additional experimental details and spectra are included in Supporting Information.

### αS Aggregation Experiments

To form pre-formed fibrils (PFFs), a 100 μM stock of unmodified αS in 1x phosphate buffered saline (PBS) was shaken at 37°C for 3-7 days. Dried aliquots of αS-L^S^8 were dissolved in PBS to 100 μM. A 1 mM stock of thioflavin-T (ThT) in 1x PBS was prepared and stored at -20 °C until use. For each percentage of thioamide αS monomer, stock solutions of thioamide-containing αS monomer, unmodified αS monomer and ThT were prepared. 50 μL of each stock solution was pipetted into a 96-well plate (Greiner Bio-One, microplate, 96 well, PS, half area, black) in triplicate. The plate was sealed and shaken in a 37 °C at 1400 rpm for 15 minutes. To make unmodified αS seeds, the PFF stock solution was diluted to 4 μM in 1x PBS. The diluted PFF stock solution was then placed in an ice water bath and sonicated at amplitude 50 for 2 minutes (1s on, 1s off). 50 μL unmodified αS seeds were then added to the well plate to give final concentrations of 2 μM seeds, 20 μM αS monomer and 5 μM ThT. Well plate was sealed tightly and shaken at 37 °C at 1400 rpm. ThT fluorescence emission was monitored using a Spark Mulitmode Microplate Reader (Tecan Trading AG; Switzerland) at 485 nm (excitation at 440nm) every few hours until aggregation was complete. Additional details and primary fluorescence data for each condition are given in Supporting Information.

Following conclusion of ThT aggregation kinetics assays, all samples were transferred to microcentrifuge tubes and pelleted (13,200 rpm, 90 min, 4 °C). The supernatant was carefully removed from each sample and resulting pellets were resuspended in 1x PBS (same volume as supernatant). 10 µL from each sample was removed and added to 2 µL of 150 µM sodium dodecyl sulfate (SDS) in water and placed on a heat block (100 °C, 20 min). After cooling samples for 5 min on ice, 6 µL 4x LDS gel loading dye was added to each sample before they were loaded onto the gel. In the meantime, leftover monomeric αS from the start of the aggregation kinetics assay was serially diluted to give 10 µL stock concentrations of 20 µM, 15 µM, 10 µM, 5 µM and 2.5 µM. Following the addition of 2 µL of 150 µM SDS in water to each standard stock, samples were boiled for 10 min and then loaded onto the gel. Gel images and analysis are shown in Supporting Information.

## Results and Discussion

### Knorr Pyrazole Activation of GB1 N-terminal Fragment

To explore the efficacy of the Knorr pyrazole thioester activation method in thioamide NCL, we synthesized GB1-L^S^_5_K_13_A (using an alanine mutant of GB1 as a pseudo-wild type protein as previously described).^41^ GB1-L^S^_5_K_13_A was synthesized from the N-terminal fragment GB1_1-12_-L^S^_5_ and the C-terminal fragment GB1_13-56_-K_13_C. The N-terminal fragment GB1_1-12_-L^S^_5_ was synthesized as an acyl hydrazide (GB1_1-12_-L^S^_5_-NHNH_2_) on 2-chlorotrityl resin and the C-terminal fragment GB1_13-56_-K_13_C was expressed as a fusion protein in *E. coli*, cleaved, and purified for NCL, both as previously reported.^41^ Reactions were performed at a 5- to 10-fold scale-up in comparison to previous ligations for incorporation of thioamides into GB1.^30, 41, 42^ Following acidic activation with acetyl acetone in the presence of mercaptophenylacetic acid (MPAA), GB1_1-12_-L^S^_5_-NHNH_2_ is fully converted to the C-terminal thioester (GB1_1-12_-L^S^_5_-SR, **1**), based matrix assisted laser desorption ionization mass spectrometry (MALDI MS) monitoring (**Figure S6**). After addition to the N-terminal Cys fragment (GB1_13-56_-K_13_C, **2**), the product (GB1-L^S^_5_K_13_C, **3**) is observed via MALDI MS after only 30 minutes (**Figure 2**). Following overnight reaction, the C-terminal thioester fragment **1** appears consumed in analytical reverse phase high performance liquid chromatography (RP-HPLC) trace. According to analytical RP-HPLC monitoring, we achieved >92% yield based on the conversion of GB1_1-12_-L^S^_5_-SR (**1**), the limiting reagent (**Figure 2** and **Figure S7**). We obtained GB1-L^S^_5_K_13_C (**3**) in 38% isolated yield (**Figure S8**).

**Figure 2.**
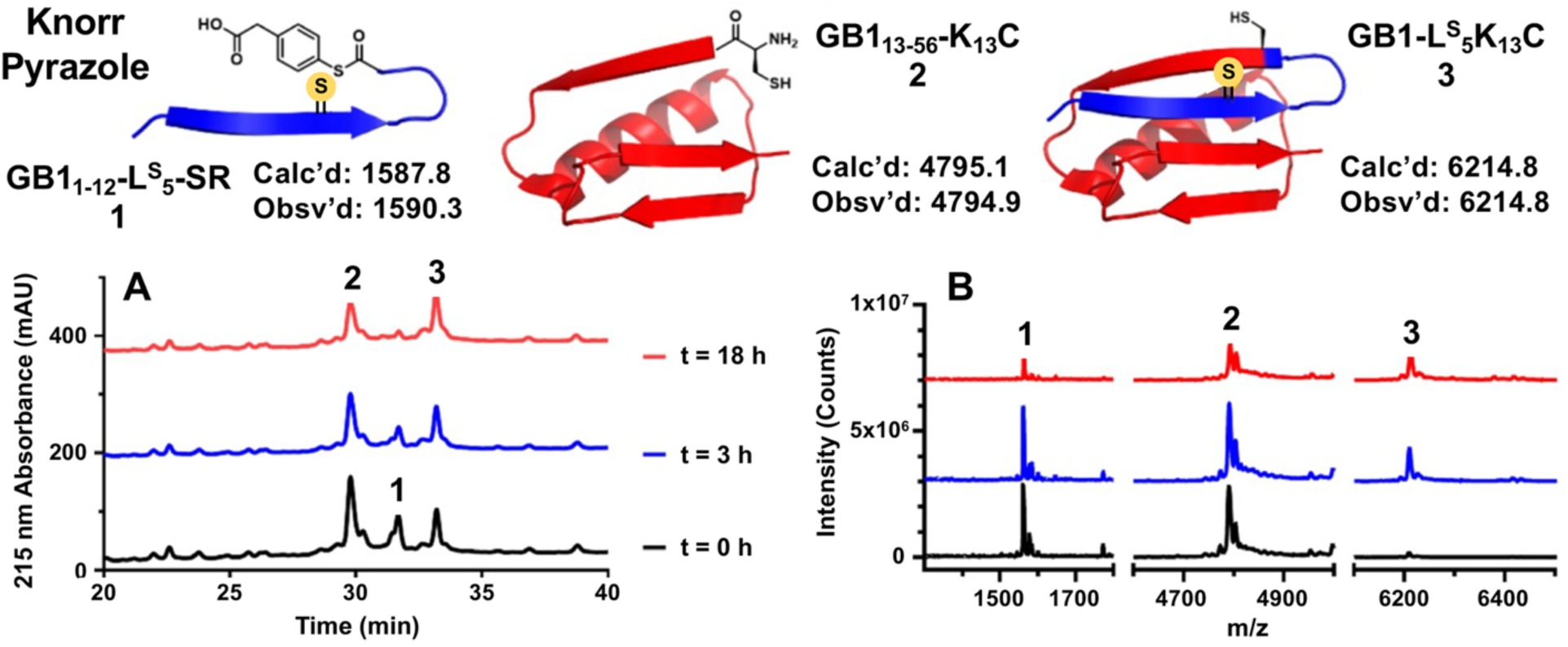
Knorr pyrazole thioester activation and NCL to semi-synthesize GB1-L^S^_5_K_13_C. The reaction was performed on the 0.71 μmol scale. Top: Schematics and calculated and observed masses for: GB1_1-12_-L^S^_5_-SR (**1**), GB1_13-56_-K_13_C (**2**), and GB1-L^S^_5_K_13_C (**3**). The analytical RP-HPLC (A) and MALDI MS (B) data demonstrate 92% conversion to NCL product GB1-L^S^_5_K_13_C (**3**) based on integration of GB1_1-12_L^S^_5_-SR (**1**) peaks.

To enable direct comparison with acyl azide activation, we performed the corresponding reactions using sodium nitrite to generate GB1_1-12_-L^S^_5_-SR (**1**) from the same GB1_1-12_-L^S^_5_-NHNH_2_ stock used in the Knorr pyrazole conversion. Activation to form the thioester was performed in 6 M guanidine buffer, pH 3, at -15 °C, with warming to room temperature and pH adjustment to ∼7 after addition of the GB1_13-56_-K_13_C fragment. Like the Knorr pyrazole ligation, ligation using acyl azide thioester activation went to >95% completion with respect to GB1_1-12_-L^S^_5_-SR. Direct comparison of the reactions shows NCL reactions using both methods proceed with essentially no side-products and >90% conversion of GB1_13-56_-K_13_C (**Figure S7**). Isolated yields ranged from 38-55% (**Table S3**).

### Knorr Pyrazole and Acyl Azide Activation of αS-L^S^_8_A_19_C

Given our success with GB1, we wished to compare the Dawson and Liu thioester activation methods using a different, larger protein construct, so the two methods were utilized in the NCL of αS with a thioamide substitution at position 8 (αS-L^S^_8_). The C-terminal fragment, αS_1-18_-L^S^_8_-NHNH_2_, was synthesized on 2-chlorotrityl resin. Upon acidic activation with acetyl acetone in the presence of MPAA, the thioamide-containing αS_1-18_-L^S^_8_-NHNH_2_ is cleanly converted to the C-terminal thioester (αS_1-18_-L^S^_8_-SR, **4**), according to MALDI MS tracking (**Figure S6**). After addition of the N-terminal cysteine-containing αS_19-140_-A_19_C (**5**) protein fragment, a small amount of NCL product (**6**) was observed after only 30 minutes (**Figure 3**). After overnight reaction, thioamide-containing thioester fragment **4** is nearly fully consumed as determined by analytical RP-HPLC. Once again, the corresponding NCL reaction was carried out using the acyl azide activation method.^43^ With either acyl azide or Knorr pyrazole thioester activation methods, some NCL product is seen after only 30 minutes, and the reaction proceeds with consumption of the thioamide-containing thioester fragment **4** overnight. When the reaction is monitored via analytical RP-HPLC, >99% of αS_1-18_-L^S^_8_-SR (**4**) is consumed during NCL with either acyl azide or Knorr pyrazole thioester activations. Both strategies afforded ligated αS-L^S^_8_A_19_C (**6**) with similar reaction rates and ∼60% conversion of αS_19-140_-A_19_C, and with no apparent side-reactions (**Figure 3**, **Figure S8**). Unfortunately, isolated yields were only ∼10% (**Table S3**), which we attribute to αS-specific losses during purification, since much higher yields were obtained for GB1.

**Figure 3.**
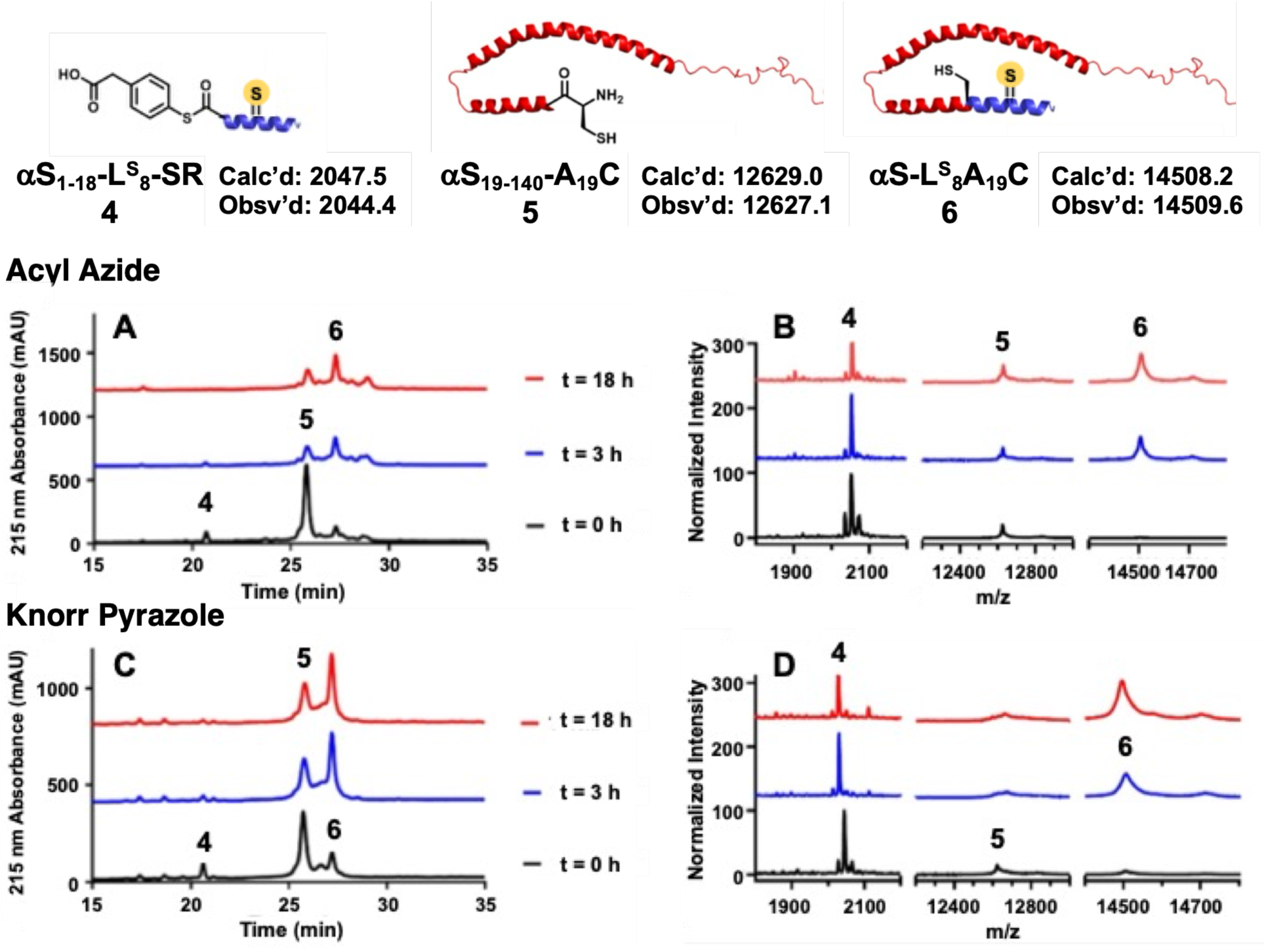
Comparison of acyl azide and Knorr pyrazole thioester activation in the semi-synthesize αS-L^S^_8_A_19_C. Top: Schematics and calculated and observed masses for the NCL constructs: αS_1-18_-L^S^_8_-SR (**4**), αS_19-140_-A_19_C (**5**) and αS-L^S^_8_A_19_C (**6**). The reaction was performed on a 1.2 μmol scale. The analytical RP-HPLC (A, C) and MALDI MS (B, D) data demonstrate conversion to NCL product **6** based on integration of the thioester (**4**) peak.

### Desulfurization of Ligation Site Cysteine

We have previously optimized methods for desulfurization of ligation site cysteines in thioamide-containing constructs using radical initiator VA-044 with thioacetamide added to prevent small amounts of thioamide-to-oxoamide conversion which were observed in prolonged reactions.^41, 42^ However, it would be beneficial to reduce the time needed for desulfurization, particularly with aggregation-prone proteins like αS. Therefore, we investigated a recently reported ultrasound induced desulfurization (USID) method^45^ in comparison to our current VA-044 induced method using a thioamidated and Cys-containing peptide substrate corresponding to the N-terminus of ligated αS (**Figure S9**). The standard VA-044 method achieved faster desulfurization than the USID method, with reaction times of 4 hours and 7 hours, respectively. While both strategies were able to access the desired desulfurized product, the USID also introduced small quantities of thioamide desulfurized product. While additional optimization of USID may be possible, the VA-044 method remains the method of choice for syntheses of thioamide containing proteins.

The ligation site cysteines in the GB1 ligation product (**3**) and the αS ligation product (**6**) were desulfurized using VA-044, with thioacetamide added to prevent desulfurization of the protein thioamide. The conversion of GB1 NCL product GB1-L^S^_5_K_13_C (**3**) proceeded cleanly in 3 hours, and the resulting GB1-L^S^_5_K_13_A protein (**3’**) was purified via RP-HPLC in a 56% yield (**Figure 4, Figure S10**, **Table S3**). The desulfurization of the cysteine-containing αS_1-140_-L^S^_8_A_19_C construct (**6**) was incomplete after 3 hours and was thus allowed to proceed overnight. Following overnight reaction, the desulfurization to αS-L^S^_8_ (**6’**) was complete and the product was purified via RP-HPLC in a 25% yield (**Figure 4, Figure S9**, **Table S3**).

**Figure 4.**
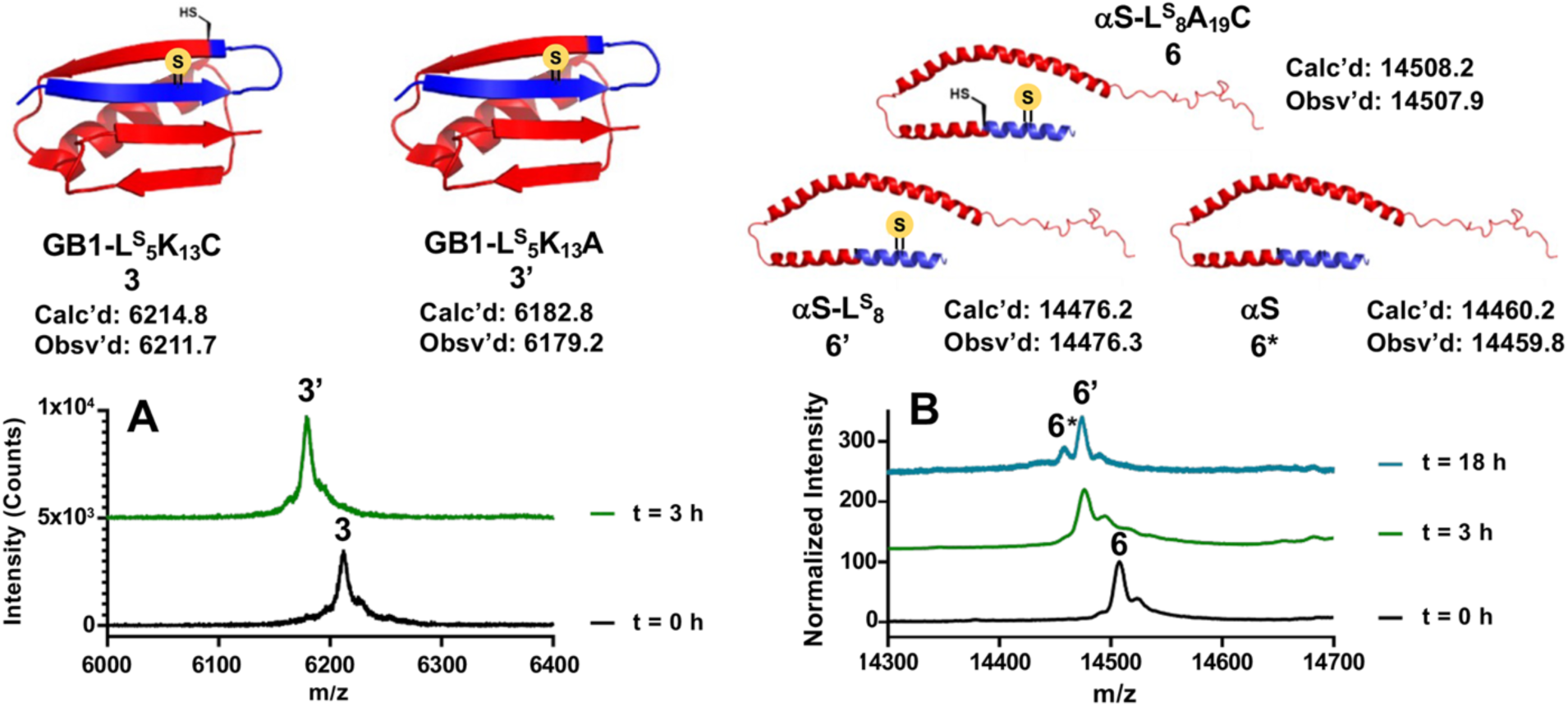
Desulfurization of ligation site cysteine to form GB1-L^S^_5_K_13_A (Left) and αS-L^S^_8_ (Right). Top: Schematics and calculated and observed masses for the Cys constructs and products: GB1-L^S^_5_K_13_C (**3**) and GB1-L^S^_5_K_13_A (**3’**); αS-L^S^_8_A_19_C (**6**), αS-L^S^_8_ (**6’**), and αS (**6\***), the thioamide desulfurization byproduct. MALDI spectra at varying timepoints are shown for the GB1-L^S^_5_K_13_C reaction (A) and αS-L^S^_8_A_19_C reaction (B).

### Biophysical Characterization of GB1

The desulfurized products from both GB1 ligations were combined for thermodynamic investigations with CD and structural studies. CD wavelength scans and thermal melts were performed with 25 µM concentrations of the protein in 20 mM Na_2_HPO_4_, pH 7.0 buffer. In CD wavelength scans, a reduction in the negative Cotton effect at 222 nm can be observed, which would could indicate destabilization of the α-helix in GB1 (**Figure S11**). A trough at 270 nm is also discernable, attributable to the thioamide itself. In CD thermal denaturation experiments, the curves can be fit to two state unfolding models to extract melting temperatures (T_m_), and GB1-L^S^_5_K_13_A (T_m_ = 60.7±0.1 °C) was found to be less thermostable than the corresponding pseudo-wild type GB1-K_13_A protein (T_m_ = 68.2±0.3 °C) by 7.5 °C (**Figure 5** and **Figure S12**). The change in free energy of unfolding (ΔΔG_U_) calculated from the T_m_ difference is -1.6 kcal mol^-1^, showing significant destabilization, consistent with our previous studies of thioamidation in WT GB1.^30^ Indeed, the L_5_ position was chosen for thioamidation in this study because it was the most perturbing position identified in those experiments. In crystal structures of WT GB1, L_5_ is found in the middle of the β-sheet portion of the protein, making both hydrogen bond donor and acceptor interactions (**Figure 5**). Given its positioning and destabilizing effects, we were intrigued to examine the structural impacts of the L_5_ thioamide.

**Figure 5.**
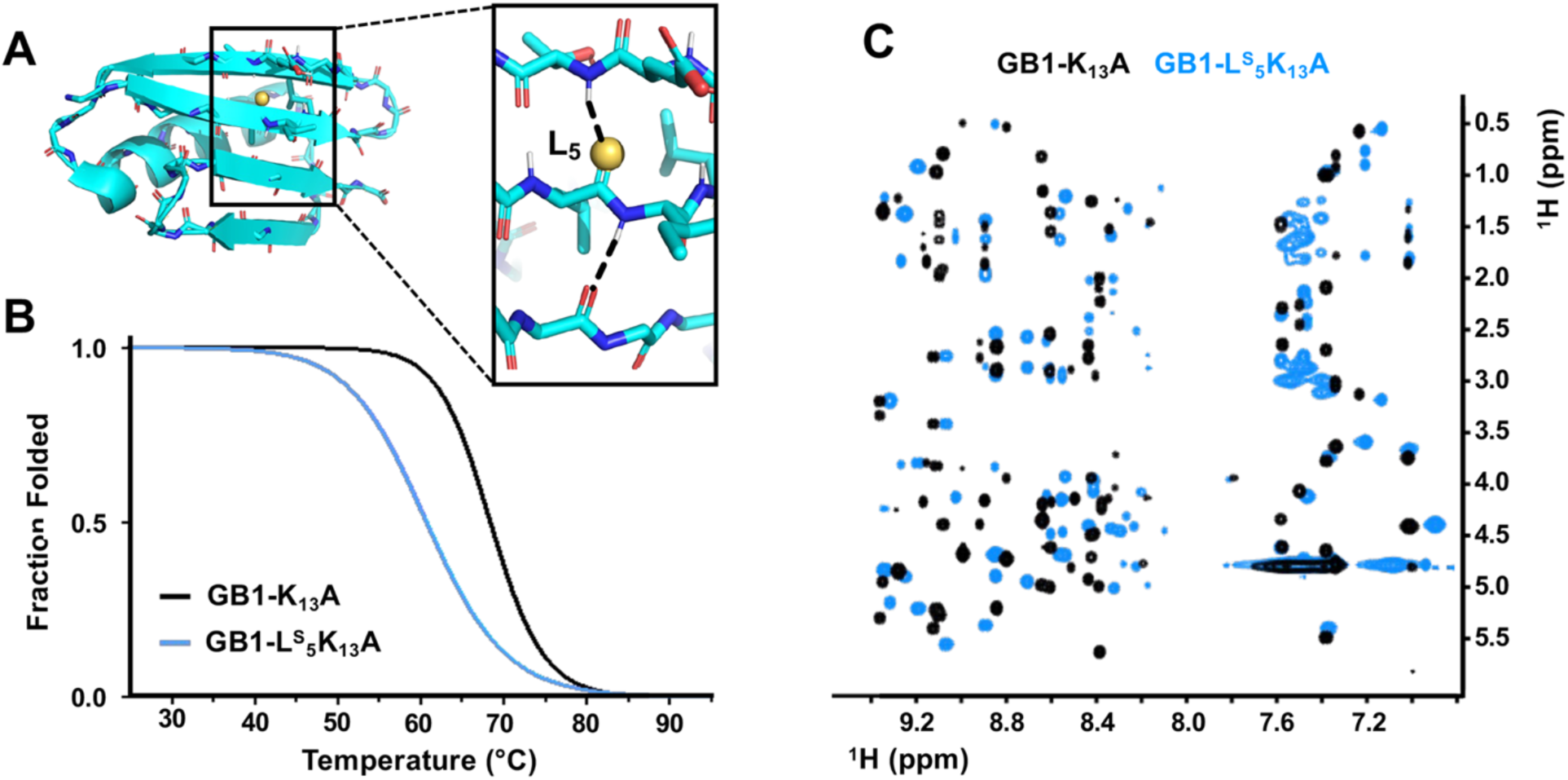
Thermodynamic and structural impact of thioamide incorporation at L_5_ in GB1-K_13_A. (A) WT GB1 structure (PDB ID 2QMT) with L_5_ H-bonding interactions highlighted in inset. (B) CD analysis demonstrates that thioamide incorporation at L_5_ results in a 7.5 °C decrease in melting temperature. (C) ^1^H-^1^H TOCSY of GB1-K_13_A (black) overlaid with GB1-L^S^_5_K_13_A (light blue) highlighting the differences in the fingerprint region (NH, Hα), located between 8.0-9.4 ppm and 3.0-6.0 ppm. Cross-peaks between 7.0-7.6 ppm are most likely from amine containing side-chains (NH_2_, Hα/ Hβ/ etc.). Cross-peaks between 8.0-9.4 ppm and 0-3.0 ppm correspond to side-chain protons (Hβ/ Hγ etc.).

We first attempted to study GB1-L^S^_5_K_13_A through X-ray crystallography. While we were able to obtain diffraction-quality crystals of the parent construct, GB1-K_13_A, and solve a structure to 1.37 Å resolution (**Figure S13**), we were not able to crystallize GB1-L^S^_5_K_13_A under similar conditions or through standard screening protocols. Therefore, we turned to NMR to gain structural insight. Since we incorporate the thioamide modification with peptide synthesis, it is prohibitively costly to acquire thioamide-containing protein that is fully isotopically labelled. Homonuclear NMR methods require large quantities of protein (100-300 µL of ∼1 mM protein solutions),^46^ but these are now accessible with the larger scale syntheses of GB1-L^S^_5_K_13_A generated here. We performed ^1^H-^1^H TOCSY experiments on 800 µM solutions of both GB1 constructs and observed dramatic differences in their spectra (**Figure S14**). In particular, the cross peaks in the fingerprint region (NH, Hα) of GB1-L^S^_5_K_13_A do not overlay with those of the parent GB1-K_13_A construct (**Figure 5**). This indicates significant disruption of the GB1 fold, consistent with the loss of stability observed by CD and with our difficulty in crystallizing GB1-L^S^_5_K_13_A. Much more modest changes to NMR spectra and ΔΔG_U_ were observed for thioamide variants of a two-stranded β-hairpin.^29^ This comparison implies that the O-to-S substitution is more disruptive in a cooperative hydrogen bonding network like the four-stranded GB1 β-sheet, where L^S^_5_ acts as both donor and acceptor.

### Aggregation Kinetics of Thioamide-Containing αS

The desulfurized products from both αS ligations were combined and used to investigate the effects that thioamide backbone modification would have on the aggregation of αS. The construct was mixed with WT αS in varying ratios and aggregation was monitored via increases in the fluorescence of thioflavin T (ThT) after seeding with 10% pre-formed αS WT fibrils. Unlike previous experiments in which the addition of 5% thioamide-containing αS had no significant effect on aggregation,^26^ dramatic effects were observed at the higher percentages of αS-L^S^_8_. ThT fluorescence was reduced proportionately, and the respective times to half-maximal ThT signal (T_1/2_) for 25%, 50%, and 100% αS-L^S^_8_ were 6.9, 12.5, and 43.9 hours, compared to the WT αS T_1/2_ of 8.5 hours (**Figure 6, Figure S16**). Control experiments using thioacetamide indicated that ThT was not quenched by the thioamide (**Figure S15**), implying that the lower signal observed in the aggregation assays could be due to decreased fibril formation. However, gel analysis showed that the amounts of insoluble aggregates ultimately formed were relatively similar under all conditions (**Figure S17, Figure S18**). Taken together, these data lead us to conclude that the thioamide modified αS is forming more amorphous aggregates that either compromise ThT binding or reduce its fluorescence activation in the bound state. Unlike the anticipated destabilizing effects of L^S^_5_ substitution in GB1, the 5-fold increase in T_1/2_ and the apparent conversion to non-fibrillar aggregates for αS-L^S^_8_ are very surprising as this region is unresolved in most solid state NMR or cryo-electron microscopy (cryo-EM) studies of αS fibrils.

**Figure 6.**
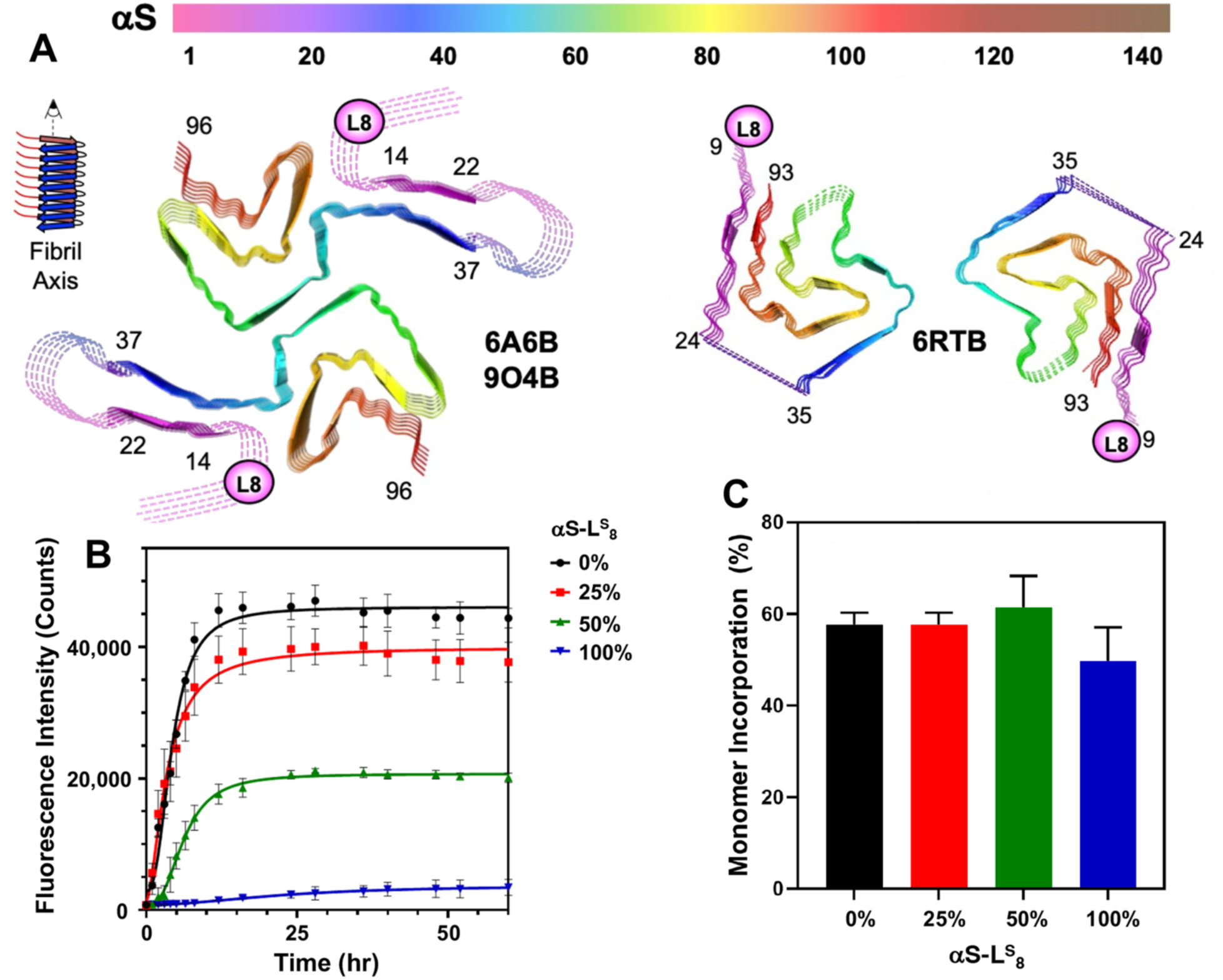
αS Aggregation Effects. (A) Structures of two canonical αS fibril polymorphs (named for the PDB IDs of representative structures) observed for WT fibrils generated under the aggregation conditions used in this study. Dashed lines indicate segments not observed in the cryo-EM structures. (B) and (C) Aggregation reactions were performed with 2 μM WT αS fibril seeds, 20 μM αS monomer (WT αS and αS-L^S^_8_ mixed in varying ratios), and 5 μM ThT, shaken at 37 °C at 1400 rpm in a 96 well plate. (B) ThT fluorescence at 485 nm measured with excitation at 440 nm. (C) Monomer incorporation into aggregates based on integration of gel bands in pelleted fraction. n=3 replicates for ThT and gel analysis.

In our studies of the effects of post-translational modifications on αS aggregation or the binding of small molecule ligands, cryo-EM structures have been solved of the WT αS fibrils prepared in our laboratory.^47, 48^ These fibrils have exhibited two conformations, fibril polymorphs that have been observed repeatedly by others. One form is exemplified by polymorph 9O4B (**Figure 6**, named for its PDB ID),^48^ a variation on the common 6A6B fold that includes a resolved N-terminal region, but only down to residue 14.^49^ The other is exemplified by 6RTB, a more rarely observed polymorph, which is resolved to residue 9.^50^ These structures, particularly 6RTB, indicate that residue 8 may indeed stack in a parallel β-sheet as part of amyloid fibril formation, where it would be expected to act as both hydrogen bond donor and acceptor. In this structural context, one can infer from our findings with GB1-L^S^_5_K_13_A that a single atom substitution is sufficient to significantly disrupt αS fibril formation, where there is tremendous cooperativity among the hydrogen bonding networks for each β-strand layer along the fibril axis. As noted above, since there is not a significant difference in the fraction of protein found in the pelleted fraction, even for 100% αS-L^S^_8_ which shows almost no ThT fluorescence, we believe that the thioamide modified αS is forming more amorphous aggregates. Further studies with other thioamide αS constructs and additional techniques such as transmission electron microscopy will be performed to better understand these effects.

The dramatic effect that a single-atom substitution has on the rate of αS fibril formation can be viewed in the context of studies that have investigated the effects of αS N-terminal mutations as well as studies of related backbone modifications in amyloid-β (Aβ) peptide aggregation. Mutations and truncations of the αS N-terminus have been shown to have significant effects on the rate of fibril formation relative to unmodified αS, highlighting the sensitivity of this seemingly disordered region to even subtle modifications.^51, 52^ The effect that backbone modification has on fibril formation has also been investigated in the context of Aβ aggregation by Kelly and coworkers. The incorporation of *E*-olefin, *N*-methyl amide, or ester isosteres into the amyloid core of Aβ_1-40_ altered both the rate and extent of aggregation due to disruption of the H-bonding network as well as steric perturbations.^53–55^ While these modifications were made in the core, other modifications, such as N-terminal truncation and pyroglutamylation of Aβ have been shown to affect aggregation despite being in regions that are not resolved in solid state NMR structures of fibrils.^56^ Taken together, these examples indicate that modifying intra- and intermolecular interactions through both backbone modification and single-residue mutations can have profound effects on aggregate formation, even when they occur outside of the ordered regions of fibrils.

## Conclusion

Studying the structural and functional differences between thioamides and oxoamides in peptides and proteins will help us to understand the role of the thioamide modification in nature and to better focus its use as a biophysical probe or peptide modifier. NCL or expressed protein ligation has been previously used to generate thioamide-containing proteins, but generally not in high enough yields for structural studies. In this work, we demonstrate that thioamide-containing peptides with C-terminal acyl hydrazides can be activated to a Knorr pyrazole with acetylacetone for generation of C-terminal thioesters for NCL. USID was also shown to be a viable alternative to VA-044 desulfurization. While the Knorr pyrazole approach did not necessarily result in higher yields of the NCL products than the corresponding acyl azide route to the thioester, we do show that both can be scaled significantly to afford sufficient quantities of GB1 and αS for studies that were previously daunting. In both cases, the resulting effects on protein folding were dramatic and generate new questions to be pursued in subsequent experiments.

We chose to study GB1-L^S^_5_K_13_A because we had observed significant destabilization for thioamide incorporation at Leu_5_ in WT GB1 and because we were successful in crystallizing the GB1-K_13_A parent construct to potentially provide a structural interpretation for destabilization. Indeed, the ΔΔG_U_ penalty of 1.6 kcal mol^-1^ for Leu_5_ thioamidation in GB1-K_13_A is less than in WT GB1 (-2.5 kcal mol^-1^).^30^ Nonetheless, GB1-L^S^_5_K_13_A proved to be unsuitable for crystallization in our hands and NMR structural characterization demonstrated that thioamide incorporation at Leu_5_ led to broad destabilization of the GB1-K_13_A structure. While this result is intriguing, structural information on thioamides at multiple positions with varying levels of perturbation needs to be acquired to gain a more holistic understanding of these effects.

The rate of aggregation of αS is significantly slowed by the incorporation of a thioamide at Leu_8_, in spite of the fact that this residue is at the edge of the ordered fibril core, or even fully disordered in all known solid state NMR and cryo-EM structures of αS fibrils.^57^ We believe that this provides evidence for a greater role of the N-terminus in fibrilization of αS, which is anticipated somewhat by observations of the effects of mutations and PTMs in this region on aggregation.^51^ Since the thioamide maintains both hydrogen bond donor and acceptor functionality, but perturbs each of them somewhat, further investigations could probe the same location with other isosteres that completely eliminate one or both of these interactions and also probe other locations in the N-terminus with thioamides and these other isosteres. These studies could have broad implications for the role of disordered and semi-ordered regions in amyloid forming proteins, complementing the studies of backbone isostere effects in the core of the Aβ peptide by Kelly and coworkers.^53, 55^

Collectively, these findings further the employment of thioamides in biophysical studies, demonstrating that dramatic destabilization can result from the perturbation of a single set of backbone interactions. However, our previous studies of GB1 have shown that other thioamide sites can have very modest influences on protein stability.^30^ Thus, the thioamide effects are clearly quite context-dependent. Learning the rules of how destabilization occurs and when comparable stabilization can be generated could guide the use thioamides in peptide and protein therapeutics and provide understanding of the role of thioamides in natural proteins. Continued improvements in synthetic methods are crucial to accessing thioamide proteins for these investigations. We will explore additional means of increasing yields for the activated protein fragments as well as their compatibility with other semi-synthesis methods that have been shown to be useful in the study of proteins like αS.^58–60^

## Supporting Information

Reaction details, analytical RP-HPLC and MALDI data on purified NCL and desulfurization constructs, expression SDS PAGE gels, supplemental analytical RP-HPLC, MALDI, CD, NMR, and aggregation kinetics data, equations, X-ray crystallographic data.

## Accession Codes

UniProtKB P37840 is the accession number for α-synuclein.

UniProtKB P06654 is the accession number for immunoglobulin protein G.

12LO is the Protein Data Bank accession code for the atomic coordinates of GB1-K_13_A.

## Acknowledgements

This work was supported by the University of Pennsylvania, the National Science Foundation (NSF CHE-2203909 to E.J.P.), and the National Institutes of Health (NIH R01-GM49758 and R35-GM163533 to D.W.C.). K.E.F. thanks NIH for funding through the Structural Biology & Molecular Biophysics Training Program (T32-GM008275). E.S.K.Y. thanks the NIH Chemistry Biology Interface Training Program (T32-GM133398) for funding. E.S.K.Y. and D.Y.F. thank the Fontaine Society for funding. A.L. thanks the Paglia Post-Bac Research Fellowship program from Carleton College for funding. The University of Pennsylvania Bruker AVANCE NEO 600 MHz NMR spectrometer was supported by NIH supplement awards R01-GM118510-03S1 and R01-GM087605-06S1, and the Vagelos Institute for Energy Science and Technology. K.E.F. acknowledges Dr. Chad Lawrence for his assistance with acquiring the NMR data. The matrix-assisted laser desorption ionization mass spectrometer was supported by NIH S10-OD030460. This research utilized the Stanford Synchrotron Radiation Lightsource (SSRL), SLAC National Accelerator Laboratory, which is supported by the U.S. Department of Energy, Office of Science, Office of Basic Energy Sciences under Contract No. DE-AC02-76SF00515. The SSRL Structural Molecular Biology Program is supported by the DOE Office of Biological and Environmental Research, and by the National Institutes of Health, National Institute of General Medical Sciences (P30-GM133894). The contents of this publication are solely the responsibility of the authors and do not necessarily represent the official views of NIGMS or NIH.

