## Supporting Information for "Improved Protein Semi-Synthesis Enables Biophysical Studies of Thioamide Destabilization of β-Sheet Interactions"

### General

Many of these methods (including cloning) are adapted from those previously described.[1]

*Reagents* *N*α-Fmoc-*N*ω-(2,2,4,6,7-pentamethyldihydro-benzofuran-5-sulfonyl)-L-arginine, 7-Azabenzotriazol-1-yloxy)tripyrrolidino-phosphonium hexafluorophosphate (PyAOP), Fmoc-Ala-OH, and Fmoc-Leu-OH were purchased from ChemImpex (Wood Dale, IL, USA). All other Fmoc-protected amino acids and resin were purchased from NovaBioChem (currently MilliporeSigma; St. Louis, MO, USA). Triisopropylsilane (TIPS) was purchased from ChemImpex or Acros (currently Fisher Scientific; Waltham, MA, USA). 1,8-Diazabicyclo[5.4.0]undec-7-ene (DBU), acetic anhydride, and thioanisole were purchased from Acros. *N*-methylmorpholine (NMM) was purchased from Acros or Alfa Aesar (currently Fisher Scientific; Waltham, MA, USA). Ethyl cyano(hydroxyimino)acetate (Oxyma) was purchased from Tokyo Chemical Industry (Tokyo, Japan). α-Cyano-4-hydroxycinnamic acid (CHCA) was purchased from Santa Cruz Biotechnology, Inc (Dallas, TX, USA) or Millipore Sigma. All other reagents and solvents were purchased from Fisher Scientific or MilliporeSigma unless otherwise specified. Milli-Q filtered (18 MΩ) water was used for all solutions. All reagents and solvents were used without further purification.

*Instrumentation* Reversed phase high performance liquid chromatography (RP-HPLC) purification was performed on an Agilent 1260 Infinity II Preparative HPLC (Santa Clara, CA, USA). RP-HPLC analytical monitoring was performed on an Agilent 1260 Infinity II Analytical HPLC (Santa Clara, CA, USA). Matrix-assisted laser desorption/ionization time-of-flight (MALDI-TOF) mass spectra were collected with a Bruker Ultraflex III or MicroFlex (Billerica, MA, USA). NMR data were acquired with a Bruker AVANCE NEO 600 MHz spectrometer. Ultraviolet-visible (UV-vis) absorption spectra were collected on a Orion AquaMate 8100 UV-vis spectrophotometer (Thermo Fisher Scientific; Waltham, MA, USA). Circular dichroism (CD) data were acquired with a Jasco J-1500 CD spectrometer. Fluorescent measurements were acquired using a Spark Mulitmode Microplate Reader (Tecan Trading AG; Switzerland).

### Fmoc-Leu$^S$-nitrobenzotriazolide Synthesis

***Scheme S1. Fmoc-Leu$^S$-Nbt (S1C) thioamide precursor synthesis***

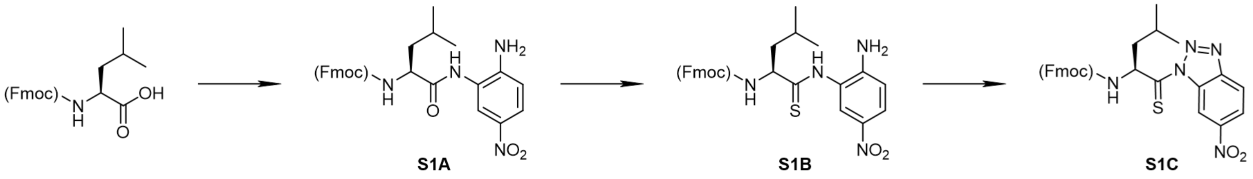


ThioLeu-Nbt (**S1C**) was synthesized as previously described[2] with the following modifications.

*Coupling of Fmoc-L Leu with 1,2-diamino-4-nitrobenzene (**S1A**)* Fmoc-Leu-OH (10 mmol, 3.53 g – 1 eq) was dissolved in 100 mL dried tetrahydrofuran (THF). NMM (20 mmol, 2.2 mL - 2 eq) was added and the reaction was cooled to -10 °C while purging with argon. Isobutyl chloroformate (IBCF, 10 mmol, 1.3 mL - 1 eq) was added dropwise to the stirred reaction. The syringe was rinsed with salt water from the cooling bath to inactivate the residual IBCF. After 15 minutes stirring at -10 °C, 4-nitro-o-phenylenediamine (10 mmol, 1.53 g – 1 eq) was added to the reaction. The reaction was stirred under argon for 2 hours at -10 °C and then at room temperature overnight. After removing the solvent in vacuo, the product was dissolved in 20 mL *N,N*-dimethylformamide (DMF) and precipitated with the addition of 1:1 saturated KCl/MilliQ $H_2O$ (500 mL). The precipitate was filtered and washed extensively with water. The residual water was removed overnight under high vacuum.

*Preparation of Fmoc L-amino ThioLeu nitroanilide (**S1B**)* Phosphorous pentasulfide ($P_4S_{10}$, 3.95 mmol, 1.76 g - 0.75 eq) and anhydrous $Na_2CO_3$ (3.95 mmol, 419 mg - 0.75 eq) were added to dry THF (56 mL) under argon and left stirring for 30 minutes (or until the phosphorous pentasulfide dissolved). After which, the compound **S1A** (5.26 mmol, 2.65 g – 1 eq) was added. The reaction was purged with argon and left to stir overnight. The next day, after removing the solvent in vacuo, the solid was resuspended in ethyl acetate and filtered over a pad of Celite©. The filtrate was washed twice with 5% $NaHCO_3$ and once with brine. The organic layers were combined and dried with $MgSO_4$. After filtration, the product was dissolved in minimal dichloromethane (DCM) and purified over silica on a Biotage Isolera One system (Biotage, LLC, Charlotte, NC, USA) with ethyl acetate/n-hexanes (25-75% ethyl acetate in *n*-hexanes).

*Preparation of Fmoc-L-amino ThioLeu nitrobenzotriazolide (**S1C**)* The compound **S1B** (4.77 mmol, 2.41 g – 1 eq) was dissolved in 95% glacial acetic acid (56 mL) (v/v in Milli-Q water) and cooled to 0°C. $NaNO_2$ (7.16 mmol, 496 mg - 1.5 eq) was added slowly and the reaction was left to stir at 0 °C under atmosphere. After 30 minutes the product was precipitated with cold MilliQ $H_2O$. The filtered precipitate was lyophilized overnight and used directly for SPPS without further purification.

### Peptide Synthesis, Purification, and Characterization

*General* Peptides were synthesized using Fmoc-based solid phase peptide synthesis (SPPS) on 2-chlorotrityl chloride resin (100-200 mesh, 1% DVB) from NovaBioChem in fritted syringes. All peptides were synthesized on either 100 or 200 µmol scales.

The resin was swelled for 30-45 minutes in DMF or DCM. A 5mL solution containing 5 eq of the first residue (Fmoc-carbazate) with 10 eq of *N,N*-diisopropylethylamine (Hünig's base) in DMF was manually coupled to resin for 30 mins. After washing the vessel with DMF and DCM, the coupling was repeated. Unreacted termini on the resin were capped by treatment with a 5mL solution of DCM/MeOH/Hünig's base (17:2:1 v/v) for 30 minutes. After draining and rinsing with DMF and DCM, the Fmoc protecting group was removed with 20% v/v piperidine in DMF (4 or 8 mL) for 2 x 10 minutes. After the first deprotection, the resin was washed with Wash 1 (4 or 8 mL each: DMF x 2, DCM, DMF). After the final deprotection, the resin was washed with Wash 2 (4 or 8mL each: (DMF, DCM) x 3, DMF).

The remaining residues up until the thioamide were coupled either manually or using a Biotage Initiator+ Alstra (Charlotte, NC, USA) automatic peptide synthesizer. Manual couplings were performed by adding 5 equivalents of the amino acid with 5 equivalents of PyAOP and 10 equivalents of Hünig's base in DMF (2 or 4 mL) to the resin. Following stirring at room temperature for 30 minutes and washing with DMF (4 or 8 mL), the coupling was repeated. Deprotections were repeated using 20% v/v piperidine in DMF as previously described. Automatic couplings were performed twice at 75°C for 5 minutes using 5 equivalents each of the amino acid, *N,N'*-diisopropylcarbodiimide (DIC), and Oxyma. Between couplings, the Fmoc protecting group was removed using 2 or 4 mL of 2% v/v DBU in DMF for 3 x 2 minutes, with DMF washes in between.

*Thioamide Coupling and Deprotection* The synthesized nitrobenzotriazolide thioamide precursor (3 eq) was dissolved in dry DMC (2-4mL) with 4 eq of Hünig's base. The mixture was added to the peptidyl-resin to stir at room temperature for 1 hour. After Wash 1, the coupling was repeated. The remaining unreacted termini were acetyl capped by treatment with 5 mL capping solution (8.4 mL DMF, 1.0 mL acetic anhydride, 0.6 mL NMM) for 2 x 10 minutes, with a DMF wash in between. Following Wash 2, the Fmoc group was removed with 2 or 4 mL of 2% v/v DBU in DMF for 3 x 2 minutes. After the first two DBU deprotections, Wash 1 was performed. After the last deprotection, Wash 2 was performed.

The remaining amino acids were coupled manually or automatically as previously described; however, all subsequent deprotections in thioamide-containing peptides were performed with 2 or 4 mL of 2% v/v DBU in DMF for 3 x 2 minutes to avoid thioamide residue epimerization and previously published side-reactions with piperidine.[3]

*Cleavage* After the desired peptide sequence was synthesized, the resin was dried extensively with DCM. Peptides were cleaved from the resin with 90% trifluoroacetic acid (TFA), 5% thioanisole, 2.5% TIPS, and 2.5% $H_2O$ for 1 hour. After removal of the solvent, the peptides were precipitated with two cold diethyl ether washes. Ether was decanted and residual ether was allowed to evaporate.

The precipitated crude peptide was then dissolved in a minimal volume of 5% (v/v) acetonitrile + 0.1% TFA in $H_2O$ + 0.1% TFA for analytical confirmation using MALDI-TOF mass spectrometry with CHCA matrix. Following positive identification of the desired peptide, the mixture was lyophilized (Labconoco; Kansas City, MO) to ensure complete removal of cleavage solvents.

*Purification* All crude peptides were purified by RP-HPLC. All purified peptides were ≥ 85% pure (based on analytical 215 nm AUC integration). Following purification, peptides were quantified by thioamide UV absorption (ε274 nm =10,169 $M^{-1}$ $cm^{-1}$).[4]

Peptides were purified on a Luna Omega PS C18 prepative column (5 μm particle size, 250 mm length, 21.2 mm diameter) (Phenomenex; Torrance, CA) with a gradient of Solvent B (acetonitrile + 0.1% TFA) in Solvent A ($H_2O$ + 0.1% TFA) ($GB1_{1-12}$-$L^{s}_{5}$-$NHNH_2$: 20-40% B, **Table S2, A**; $\alpha S_{1-18}$-$L^{s}_{8}$-$NHNH_2$: 20-35% B, **Table S2, B**). $GB1_{1-12}$-$L^{s}_{5}$-$NHNH_2$ was subjected to second-pass RP-HPLC purification on a Luna Omega PS C18 prepative column (5 μm particle size, 250 mm length, 21.2 mm diameter) with a 23-27% gradient (**Table S2**, **C**). Following identification of the peptide with MALDI-TOF-MS using CHCA matrix, the peptide was lyophilized.

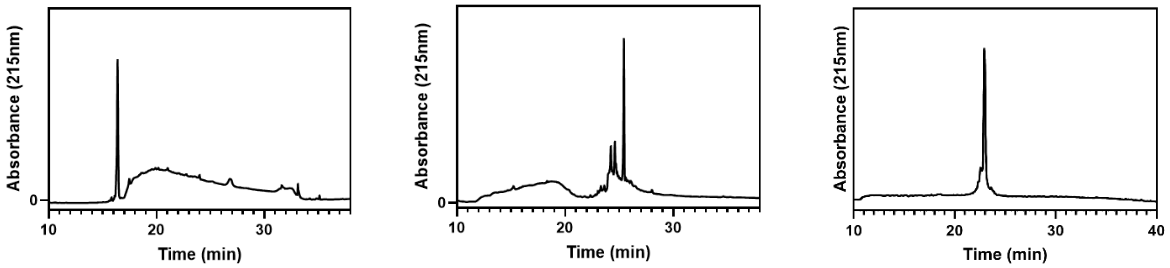


**Figure S1.** Analytical HPLC traces of synthesized GB1$_{1\text{-}12}$-L$^{s}_{5}$-NHNH$_2$ (left, Gradient D, **Table S2**), αS$_{1\text{-}18}$-L$^{s}_{8}$-NHNH$_2$ (middle, Gradient D, **Table S2**) and αS$_{1\text{-}18}$-L$^{s}_{8}$ A$_{17}$C (right, Gradient E, **Table S2**).

**Table S1.** MALDI characterization, gradient of purification, and analytical retention time of the synthesized peptides

| **Peptide** | **[M+H]$^+$ (m/z)** | | **[M+Na]$^+$ (m/z)** | | **[M+K]$^+$ (m/z)** | | **Gradient** | **Analytical Retention Time*** |
|---|---|---|---|---|---|---|---|---|
| | **Exp** | **Obs** | **Exp** | **Obs** | **Exp** | **Obs** | | |
| GB1$_{1\text{-}12}$-L$^{s}_{5}$--NHNH$_2$ | 1451.86 | 1451.54 | 1474.95 | --- | 1491.06 | --- | **A, C** | 24.169 |
| αS$_{1\text{-}18}$-L$^{s}_{8}$--NHNH$_2$ | 1911.38 | 1911.78 | 1934.37 | --- | 1950.48 | -- | **B** | 25.425 |
| αS$_{1\text{-}18}$-L$^{s}_{8}$ A$_{17}$C | 1929.41 | 1929.23 | 1952.41 | 1951.21 | 1969.41 | -- | **E** | 22.921 |

**Table S2.** HPLC gradients for peptide purification (solvent A = Milli-Q water with 0.1% TFA, solvent B = acetonitrile with 0.1% TFA)

| **Grad.** | **Time** | **% B** | **Grad.** | **Time** | **%B** |
|---|---|---|---|---|---|
| **A** | 0.00 | 5 | **B** | 0.00 | 5 |
| | 3.00 | 5 | | 3.00 | 5 |
| | 8.00 | 20 | | 8.00 | 20 |
| | 11.00 | 20 | | 11.00 | 20 |
| | 31.00 | 40 | | 31.00 | 35 |

| | | | | | |
|---|---|---|---|---|---|
| | 34.00 | 40 | | 34.00 | 35 |
| | 38.00 | 100 | | 38.00 | 100 |
| | 43.00 | 100 | | 43.00 | 100 |
| | 45.00 | 5 | | 45.00 | 5 |
| **Grad.** | **Time** | **% B** | **Grad.** | **Time** | **%B** |
| **C** | 0.00 | 5 | **D** | 0.00 | 5 |
| | 3.00 | 5 | | 2.00 | 5 |
| | 6.00 | 23 | | 5.00 | 5 |
| | 12.00 | 23 | | 8.00 | 5 |
| | 32.00 | 27 | | 38.00 | 90 |
| | 35.00 | 35 | | 40.00 | 100 |
| | 38.00 | 100 | | 45.00 | 100 |
| | 43.00 | 100 | | 48.00 | 5 |
| | 45.00 | 5 | | | |
| **Grad.** | **Time** | **% B** | | | |
| **E** | 0.00 | 5 | | | |
| | 3.00 | 5 | | | |
| | 8.00 | 25 | | | |
| | 11.00 | 25 | | | |
| | 31.00 | 43 | | | |
| | 34.00 | 43 | | | |
| | 36.00 | 100 | | | |
| | 39.00 | 100 | | | |
| | 41.00 | 5 | | | |
| | 45.00 | 5 | | | |

### Expression and Purification

*General* Centrifugation was performed with a Sorvall RC-5 centrifuge using SS-34 and GS3 rotors (Watham, MA, USA). Cells were lysed using a Q700 sonicator from Qsonica (Newtown, CT, USA). UV-Vis absorbance and $OD_{600}$ measurements were made on a Thermo Orion AquaMate 8100 UV-Vis Spectrometer from Thermo Scientific (Waltham, MA, USA). Expressed protein purification was performed on an ÄKTA Pure 25 Fast Protein Liquid Chromatography (FPLC) (Cytiva, Marlborough, MA) using a Superdex 75 pg column (Cytiva, Marlborough, MA, USA). The dialysis tubing used was made of regenerated cellulose from Spectrum Labs (Waltham, MA, USA) or Slide-A-Lyzer™ dialysis cassettes (3K MWCO) from Thermo Scientific (Waltham, MA, USA). The protein stocks were concentrated using Amicon Ultra-4 10k MWCO Spin Filters and sterile filtered with 0.22 µm PES Millex-GP Syringe Filter from EMD Millipore (Burlington, MA, USA). Isopropyl β-D-1-thiogalactopyranoside (IPTG) was purchased from LabScientific Inc. (Highlands, NJ, USA). Protease inhibitor cocktail cOmplete mini EDTA-Free tablets were purchased from Roche (Basel, Switzerland). Nickel agarose resin (high density) was purchased from GoldBio (St. Louis, MO, USA). Gels were run utilizing a SDS-PAGE apparatus from Bio-Rad (Hercules, CA, USA). The Spectra Multicolor Low Range Protein Ladder and GelCode Blue Stain Reagent was purchased from Thermo Scientific (Waltham, MA, USA). Gels were imaged using the G:Box Mini from Syngene (Frederick, MD, USA). NuPAGE™ LDS Sample Buffer (4X) was purchased using from Invitrogen/Thermo Fisher (Waltham, MA, USA).

*C-Terminal Catalytic Domain of Ulp-1* $His_6$-Ulp1 was expressed in E. coli BL21(DE3) cells based on the previously published procedure.[5] The plasmid pFGET19_Ulp1 was generated by Hideo Iwai (Addgene plasmid #64697). After growth and protein induction, cells were resuspended in 40 mM Tris, 500 mM NaCl, 1 mM phenylmethylsulfonyl fluoride (PMSF), pH 8.3. The cells were lysed by sonication and the supernatant of the cell lysate was applied to nickel agarose resin equilibrated with 50 mM 4-(2-hydroxyethyl)-1-piperazineethanesulfonic acid (HEPES), 300 mM NaCl, 10 mM imidazole pH 7.5. The protein was allowed to incubate with the nickel resin for 1 hour at 4 ˚C with mixing. The resin was washed with the equilibration buffer. The protein was eluted with the equilibration buffer with 300 mM imidazole. The protein was dialyzed into 20 mM Tris, 150 mM NaCl pH 8.0. The material was spin concentrated to less than 10 mL and purified over two runs on a Superdex 75 pg column with an isocratic elution of 15% 20 mM Tris, 150 mM NaCl pH 8.0 over 1.2 column volumes (**Table S4, A**). The desired protein eluted after 0.55 column volume (70 min). The material was concentrated to 1 mL and 20 µL of freshly prepared dithiothreitol (DTT) and 980 µL 50% (v/v) glycerol in water were added. The protein was aliquoted and stored at -80 ˚C until needed. The molecular weight of the product is 27,394.29 g/mol.

*His$_6$-SUMO-GB1$_{13\text{-}56}$ K$_{13}$C* His$_6$-SUMO-GB1$_{13\text{-}56}$-K$_{13}$C was expressed in *E. coli* BL21(DE3) cells. The cells were resuspended in 40 mM Tris, 1 mM PMSF, pH 8.3 with a Roche protease inhibitor cocktail tablet (EDTA-free). The cells were lysed by sonication and the supernatant of the cell lysate was applied to nickel agarose resin equilibrated with 50 mM HEPES pH 7.5. The protein was allowed to incubate with the nickel resin for 1 hour at 4 ˚C with mixing. The resin was washed as follows: equilibration buffer and equilibration buffer with 5 mM imidazole. The protein was eluted with the equilibration buffer with 300 mM imidazole. To the eluted protein, 24 μL Ulp-1 and freshly prepared DTT to a final concentration of 5 mM were added. Removal of the His$_6$-SUMO tag was achieved by mixing at 4 ˚C for 24 hours. The DTT was removed by dialysis into 20 mM Tris pH 8.0 with 500-1000 Da MWCO dialysis tubing. The sample was incubated with Ni resin equilibrated with dialysis buffer for 1 hour at 4 ˚C with mixing. The desired protein was collected in the flow-though. After dialysis into 20 mM Tris pH 8.0 (with 500-1000 Da MWCO dialysis tubing), the protein was purified via anion-exchange chromatography with 0-40% gradient (over 40 CV) with 1 M NaCl as the elutant (**Table S*4*, B**). The desired fractions were identified with MALDI-TOF MS using CHCA matrix. The product elutes at 26% B with a molecular weight of 4794.06 g/mol. Addition of tris(2-carboxyethyl)phosphine hydrochloride (TCEP) is necessary prior to purification to reduce any disulfide dimers of the N-terminal cysteine. The protein was dialyzed into Milli-Q water (with 500-1000 Da MWCO dialysis tubing), quantified by UV ($\varepsilon_{280nm}$ = 8,480 $M^{-1}$ $cm^{-1}$), aliquoted and lyophilized. Additional details can be found as in previous reports.[1]

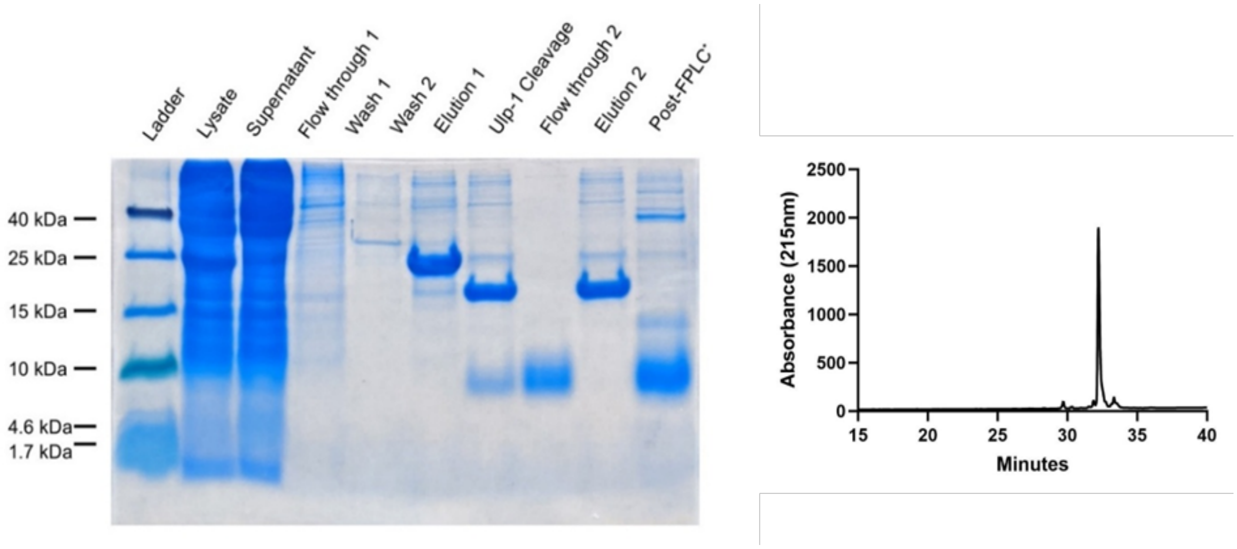


**Figure S2.** 14% Tris-Tricine SDS-PAGE gel of GB1$_{13\text{-}56}$-K$_{13}$C expression and purification and analytical HPLC post-purification (**Table S4, C**). Gel was stained with Coomassie sensitive stain (50 g aluminum sulfate, 100 mL ethanol, 200 mg Coomassie Brilliant Blue G-250, 23.5 mL O-phosphoric acid and 800+ mL of Milli-Q water for a final volume of 1000 mL). The aggregates observed are a product of lyophilization.

*GB1 K$_{13}$A-NpuDnaE-His$_6$* The pTXB1 GB1 K$_{13}$A-NpuDnaE-His$_6$ plasmid was transformed into BL21(DE3) *E. coli* cells by heat-shock and grown on LB-agar plates with ampicillin (100 μL/mL). Plates were incubated at 37 °C for at least 16 hours. Individual colonies were isolated and grown in 5 mL sterile LB media with ampicillin (100 μL/mL) and grown overnight at 37 °C with 250 RPM shaking. To each liter of sterile LB media up to three primary cultures were added and grown until saturation at 37 °C with 250 RPM shaking. Once an $OD_{600}$ of 0.6-0.8 was reached, the cells were induced with IPTG to a final concentration of 1 mM. The cells were incubated at 18 °C overnight with 250 RPM shaking. The cells were harvested the following day by centrifugation at 4,000 RPM (2,704 x g) in a GS3 rotor with a Sorvall RC-5 centrifuge for 20 minutes at 4°C. The supernatant was discarded, and the pellet was resuspended in 30 mL of 40 mM Tris pH 8.3 with 1 mM PMSF and a broad-spectrum protease inhibitor tablet. The resuspended cells were lysed via sonication (35 amp power, 1 second pulse, 2 second rest for 5 minutes total) and then pelleted at 14,000 RPM (23,426 x g) with a SS-34 rotor in a Sorvall RC-5 centrifuge.

Nickel agarose resin was added to a fritted column for a settled volume of 5 mL. The resin was equilibrated with 25 mL of 50 mM HEPES pH 7.5. The supernatant of the cell lysate was incubated with the equilibrated resin for at least 60 minutes with rotation at 4 °C. The lysate was applied back to the fritted column and the flow-through was allowed to drain. The resin was washed with 25 mL of 50 mM HEPES pH 7.5 and 25 mL of 50 mM HEPES, 5 mM imidazole pH 7.5. The desired construct was eluted from the resin with 12 mL of 50 mM HEPES, 300 mM imidazole pH 7.5. 2-mecaptoethanol (BME) was added to a final concentration of 1 M and left to mix at 37 °C for 60 hours. The sample was transferred to 3.5 kDa MWCO dialysis tubing and was dialyzed into 20 mM Tris pH 8.0 at 4 °C. The non-His-tagged material was isolated with a second-nickel purification. Nickel agarose resin was added to a fritted column for a settled volume of 5 mL. The resin was equilibrated with 25 mL of the dialysis buffer (20 mM Tris pH 8.0). The sample was incubated with the equilibrated resin for at least 60 minutes with rotation at 4 °C. The resin was applied back to the fritted column and the flow-through containing the non-His-tagged material was collected. The flow-through was then dialyzed into 20 mM Tris pH 8.0.

The construct was further purified via anion-exchange chromatography with a 0-30% gradient (over 25 column volumes) with 1 M NaCl as the elutant (**Table S4, D**). The desired fractions were identified with MALDI-TOF MS using CHCA matrix. The product elutes at 12.5% B with a molecular weight of 6165.74 g/mol. The protein was dialyzed into Milli-Q water, quantified by UV ($\varepsilon_{280nm}$ = 9,970 $M^{-1}$ $cm^{-1}$), aliquoted and lyophilized.

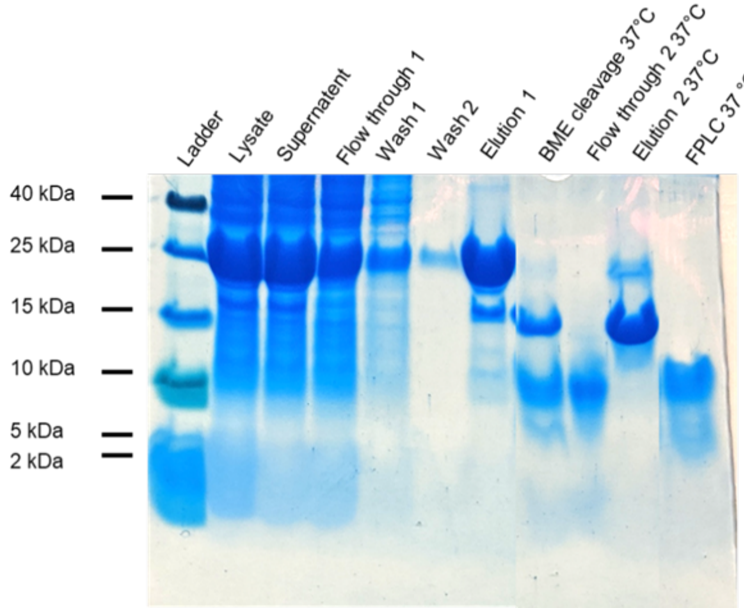

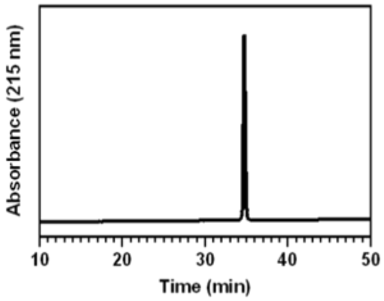


**Figure S3.** 14% Tris-Tricine SDS-PAGE gel of GB1-$K_{13}$A expression/ purification and analytical RP-HPLC post-purification (**Table S4, C**).

*His₆-GyrA-αS₁₉₋₁₄₀ A₁₉C* His$_6$-GyrA-αS$_{19-140}$-A$_{19}$C plasmid was transformed into BL21(DE3) *E. coli* cells by heat shocking at 42 °C. Cells were grown on LB-agar plates with ampicillin (100 µL/mL) overnight and single colonies were picked to inoculate primary cultures in LB media supplemented with 0.1 mg/mL ampicillin. Secondary cultures supplemented with ampicillin were incubated at 37 °C in a shaker at 250 RPM until OD reached 0.6. Expression was induced with IPTG and induced cells were grown at 18 °C in a shaker at 250 RPM overnight. Culture was centrifuged (4000 RPM, 20 min, 4 °C) and resulting cell pellets were resuspended in a 40 mM Tris pH 8.3 resuspension buffer with Roche protease inhibitor cocktail tablet (EDTA-free). Resuspended cells were lysed by sonication in an ice bath (amplitude 30, 5 min, 1 s ON, 1s OFF). The resulting lysate was centrifuged (14000 RPM, 30 min, 4 °C) and the supernatant that was applied to a Ni-NTA resin equilibrated with 50 mM HEPES, pH 7.5. The supernatant was allowed to incubate at 4 °C with the Ni-NTA resin for at least 1 hour with mixing. Following resin washes with 50 mM HEPES pH 7.5 and 50 mM HEPES, 5 mM imidazole pH 7.5, the protein of interest was eluted from the resin with a 50 mM, 300 mM imidazole pH 7.5 buffer. Intein cleavage was carried out by incubation with 200 mM BME on a rotisserie overnight at room temperature. The cleaved protein of interest was dialyzed (3 kDa MWCO tubing) into 20 mM Tris, pH 8.0 overnight. Dialyzed protein of interest was allowed to incubate with 50 mM HEPES washed Ni-NTA resin for at least 1 hour, followed by a second Ni-NTA column purification to remove any free intein. Methoxyamine hydrochloride was added to a concentration of 200 mM, and the reaction was incubated at 37 °C for 4 hours and monitored by MALDI-TOF MS. TCEP was added to a final concentration of 1mM and was left to incubate for ~10 minutes before filtering using a 22 µm syringe filter and purifying by RP-HPLC using a Jupiter C4 semi-preparative column (10 µm particle size, 250 mm length, 10 mm diameter) with a 33-43% gradient (**Table S4, E**).

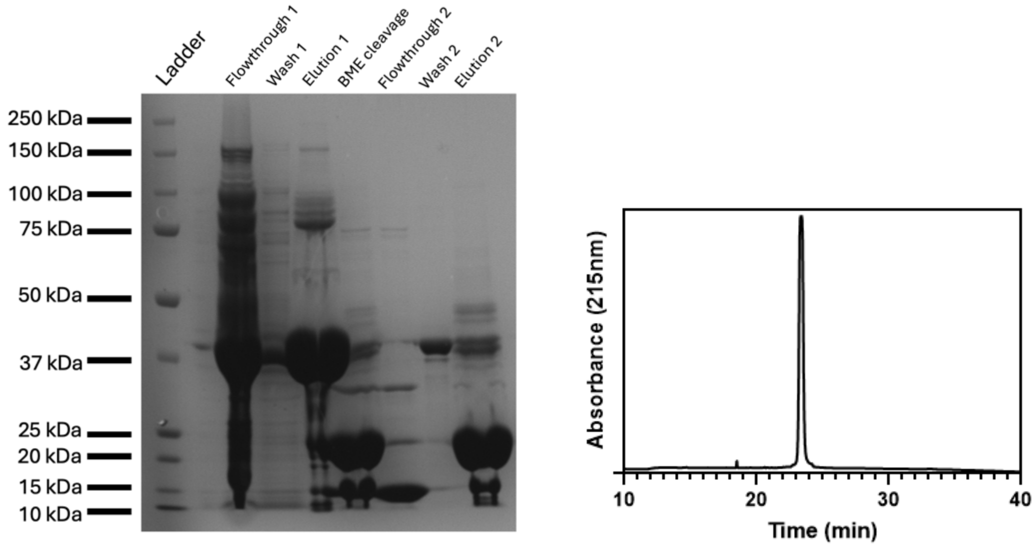


**Figure S4.** 14% Tris-Tricine SDS-PAGE gel of αS$_{19-140}$-A$_{19}$C expression and purification and analytical HPLC post-purification (**Table S4, C**). Gel was stained with Coomassie sensitive stain (50 g aluminum sulfate, 100 mL ethanol, 200 mg Coomassie Brilliant Blue G-250, 23.5 mL O-phosphoric acid and 800+ mL of Milli-Q water for a final volume of 1000 mL).

*$His_6$-GyrA-$\alpha S_{1-140}$* $His_6$-GyrA-αS plasmid was transformed into BL21(DE3) *E. coli* cells by heat shocking at 42 °C. Cells were grown on LB-agar plates with ampicillin (100 µL/mL) overnight and single colonies were picked to inoculate primary cultures in LB media supplemented with 0.1 mg/mL ampicillin. Secondary cultures supplemented with ampicillin were incubated at 37 °C in a shaker at 250 RPM until OD reached 0.6. Expression was induced with IPTG and induced cells were grown at 18 °C in a shaker at 250 RPM overnight. Culture was centrifuged (4000 RPM, 20 min, 4 °C) and resulting cell pellets were resuspended in a 40 mM Tris pH 8.3 resuspension buffer with Roche protease inhibitor cocktail tablet (EDTA-free). Resuspended cells were lysed by sonication in an ice bath (amplitude 30, 5 min, 1 s ON, 1s OFF). The resulting lysate was centrifuged (14000 rpm, 30 min, 4 °C) and the supernatant was applied to a Ni-NTA resin equilibrated with 50 mM HEPES, pH 7.5. The supernatant was allowed to incubate at 4 °C with the Ni-NTA resin for at least 1 hour with mixing. Following resin washes with 50 mM HEPES pH 7.5 and 50 mM HEPES, 5 mM imidazole pH 7.5, the protein of interest was eluted from the resin with a 50 mM, 300 mM imidazole pH 7.5 buffer. Intein cleavage was carried out by incubation with 200 mM BME on a rotisserie overnight at room temperature. Cleaved protein of interest was dialyzed (3 kDa MWCO tubing) into 20 mM Tris, pH 8.0 overnight. Dialyzed protein of interest was allowed to incubate with 50mM HEPES washed Ni-NTA resin for at least 1 hour, followed by a second Ni-NTA column purification to remove any free intein. The dialyzed protein was filtered using a 22 µM syringe filter and purified over FPLC (Cytivia, Marlborough, MA)) with a 20-35% gradient using a 5 mL Q column (**Table S4, F**). FPLC fractions containing the protein of interest were pooled, buffered exchanged into 1x PBS and spin concentrated (Amicon, 3500 kDa MWCO) to roughly 100 µM before being flash-frozen and stored at -80 °C.

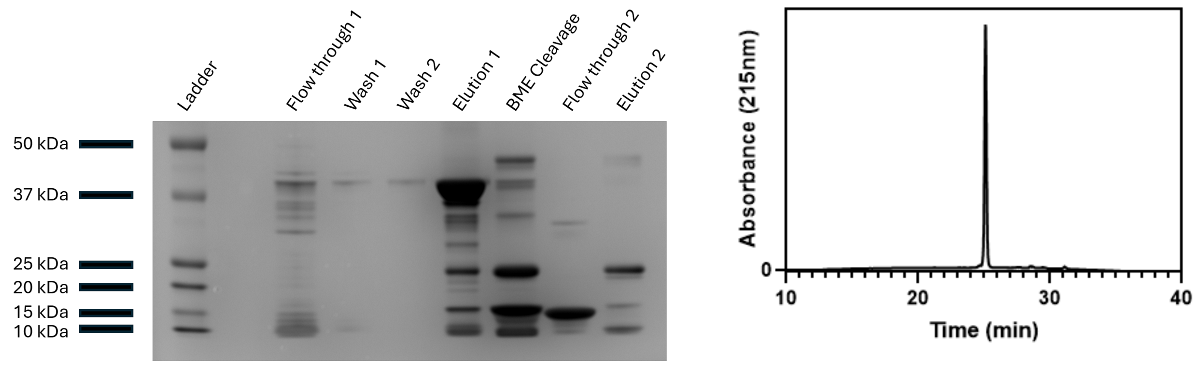


**Figure S5.** 14% Tris-Tricine SDS-PAGE gel of $\alpha S_{1-140}$ expression and purification and analytical HPLC post purification (**Table S4, C**). Gel was stained with Coomassie sensitive stain (50 g aluminum sulfate, 100 mL ethanol, 200 mg Coomassie Brilliant Blue G-250, 23.5 mL O-phosphoric acid and 800+ mL of Milli-Q water for a final volume of 1000 mL).

### Native Chemical Ligations

*Acyl Azide Ligation* The activation and native chemical ligation (NCL) buffers were made the day before and argon purged prior to starting the ligation (activation buffer: 6 M guanidine (Gdn) HCl, 0.2 M $NaH_2PO_4$ pH 3.0, NCL buffer: 6 M Gdn HCl, 0.2 M phosphate, 0.2 M 4-mercaptophenylacetic acid (MPAA) pH 7.0). The acyl hydrazide peptide was dissolved in the activation buffer to achieve a 5 mM peptide concentration. The peptide was left stirring in a -15 ˚C in an NaCl/ice bath in a Dewar bowl. After the peptide was cooled, 10 equivalents of sodium nitrate (from a 1 M stock in sterile Milli-Q water) was added and the peptide was left to stir for 15 minutes. In the meanwhile, the N-terminal cysteine containing protein was dissolved in NCL buffer to achieve a ratio of 16:1 MPAA/ $NaNO_2$. After the 15-minute incubation, the protein was added to the chilled peptide. The pH was monitored and carefully adjusted to ~ pH 7.0. The reaction was monitored by MALDI-TOF MS and analytical RP-HPLC. MALDI-TOF MS timepoint aliquots were diluted by 100x with Milli-Q water with 0.1% TFA and co-spotted with CHCA matrix. For RP-HPLC analytical monitoring, 6 µL of the reaction was mixed with 184 µL Milli-Q water + 0.1% TFA and 10 µL of 0.5 M TCEP neutral (25 mM final TCEP). After incubation for 10 minutes, the sample was injected on a C8 column (Luna C8(2) with 5 µm particle size, 150 mm length, 4.6 mm diameter, 100 Å pore size) (Phenomenex)) and analyzed with a 20-35% over 40 minutes (**Table S4**, **G**). A freshly prepared solution of TCEP neutral was added to the reaction after 3-4 hours to a final concentration of 50 mM TCEP.

The ligation was allowed to proceed at room temperature overnight. After the reaction was determined complete by MALDI-TOF MS, the NCL was slowly diluted with Milli-Q water and TCEP was added. The sample was either dialyzed into 20 mM Tris pH 8.0 or directly purified on a C18 semi-preparative column (Luna Omega PS C18 with 5 µm particle size, 250 mm length, 10 mm diameter, 100 Å pore size) (Phenomenex)) with 20-40% gradient (**Table S4, H**). The product was quantified by UV ($\varepsilon_{274nm}$ = 19,736 $M^{-1}$ $cm^{-1}$) and lyophilized. Additional details can be found as previously described.[1]

*Knorr Pyrazole Activation* The acyl hydrazide peptide was dissolved in 6 M Gdn HCl to 2 mM concentration. Solid MPAA was weighed out to achieve a 0.2 M MPAA solution and was carefully transferred to the peptide solution. From a 100 mM stock in Milli-Q water, acyl acetone (2.5 eq, 0.975 g/mL) was added to the peptide and allowed to stir at room temperature. The reaction was monitored by MALDI-TOF MS via a 100x dilution in Milli-Q water with 0.1 % TFA. After 3 hours, the thioester formation was determined complete by analytical. The NCL buffer (6 M Gdn HCl, 0.2 M $Na_2HPO_4$ pH 8.5) was purged with argon. The N-terminal cysteine containing protein was dissolved in the NCL buffer to a concentration of 2 mM protein with a final concentration of 50 mM TCEP (from a 0.5 M TCEP neutral stock). The protein was added to the peptide and the pH was monitored and adjusted to pH 7.0-7.5. The reaction was monitored by MALDI-TOF MS as described above. For analytical RP-HPLC monitoring, 10 µL of the reaction was mixed with 170 µL Milli-Q water + 0.1% TFA and 20 µL of 0.5 M TCEP neutral (50 mM final TCEP). After incubation for 10 minutes, the sample was injected on a C8 column (Luna C8(2) with 5 µm particle size, 150 mm length, 4.6 mm diameter, 100 Å pore size) (Phenomenex)) and analyzed with a 20-35% gradient over 40 minutes (**Table S4**, **G**).

The ligation was allowed to proceed at room temperature overnight. After the reaction was determined complete by MALDI MS, the NCL was slowly diluted with Milli-Q water and TCEP was added. The sample was either dialyzed into 20 mM Tris, pH 8.0 or directly purified on a C18 semi-preparative column (Luna Omega PS C18 with 5 µm particle size, 250 mm length, 10 mm diameter, 100 Å pore size) (Phenomenex)) with 20-40% gradient (**Table S4**, **H**). The product was quantified by UV ($\varepsilon_{274nm}$ = 19,736 $M^{-1}$ $cm^{-1}$) and lyophilized.

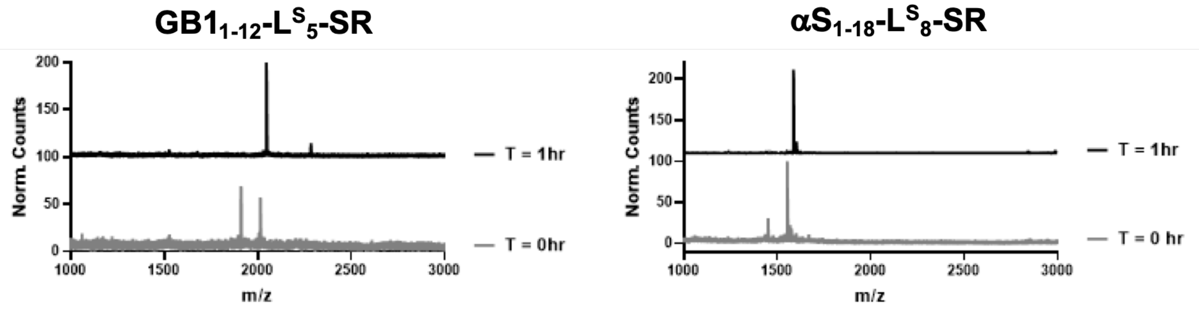


**Figure S6**. Thioester conversion of αS-L$^S_8$-SR (Left) and GB1L$^S_5$-SR (Right) using Knorr pyrazole activation of C-terminal acyl hydrazide. MALDI data demonstrates efficient thioester activation after 1 hr. Calculated masses are as follows: αS-L$^S_8$-NHNH$_2$ $[M+H]^+$ 1911.38, αS-L$^S_8$-SR $[M+H]^+$ 2047.54, GB1-L$^S_5$-NHNH$_2$ $[M+H]^+$ 1451.86, GB1-L$^S_5$-SR $[M+H]^+$ 1588.02. Observed masses are as follows: αS-L$^S_8$-SR (T = 0 hr: 1910.1, 2014.3, T = 3 hr: 2046.3), GB1-L$^S_5$-SR (T = 0 hr: 1451.6, 1553.3, T = 3 hr: 1587.6).

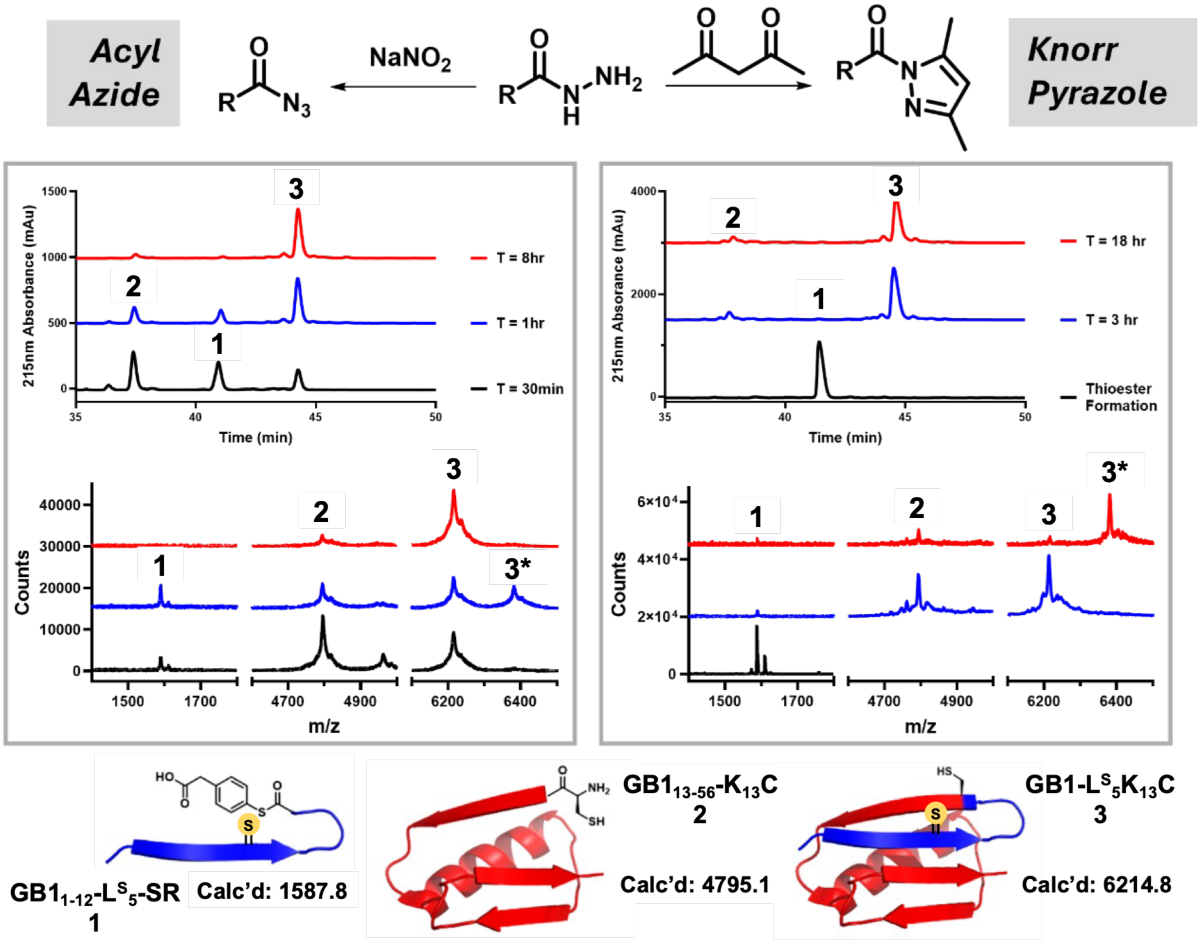


**Figure S7**. Comparison of Dawson (right) versus Liu (left) thioester activation to semi-synthesize GB1-L$^{S}_{5}$ K$_{13}$C. The Dawson activation was performed on a 0.71 μmol scale. The Liu activation was performed on a 1.0 μmol scale. The analytical (top) and MALDI (bottom) data demonstrate complete conversion to NCL product. The peak in the analytical that elutes at 27 minutes is MPAA. Below: The calculated masses for the NCL constructs: GB1$_{1-12}$-L$^{S}_{5}$-MPAA (**1**), GB1$_{13-56}$-K$_{13}$C (**2**), and GB1-L$^{S}_{5}$K$_{13}$C (**3**), MPAA disulfide adduct (**3***). (T = 0 hr: Azide 1588.9, 4795.7, 6215.6 Pyrazole 1587.7. T = 3 hr: Azide 1589.6, 4795.3, 6215.7, 6382.5 Pyrazole 1589.7, 4793.8, 6213.2. T = 18 hr: Azide 4794.9, 6216.9 Pyrazole 1589.5, 4794.3, 6216.3, 6380.3.)

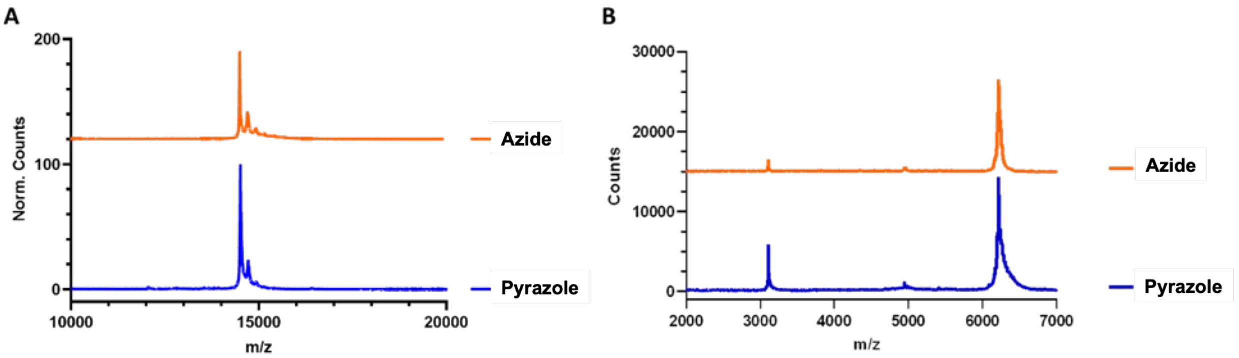


**Figure S8.** MALDI of the purified αS-$L^S_8$ $A_{19}C$ (A) and GB1-$L^S_5$ $K_{13}C$ (B) constructs. **A)** (calculated = [M+H]+ 14508.22, observed = [M+H]+ Pyrazole 14514.58, Azide 14493.09). **B)** (calculated = $[M+H]^+$ 6214.84/ $[M+2H]^{2+}$ 3107.92, observed = $[M+H]^+$ Pyrazole 6216.82, Azide 6217.15/ $[M+2H]^{2+}$ Pyrazole 3108.80, Azide 3109.35)

*Ultrasound Induced Desulfurization (USID)* Additional details can be found as previously described.[6] Ultrasound induced desulfurization was performed in a 0.5 M TCEP, 0.2 M thioacetamide pH 7.0 buffer. The buffer was prepared on the same day and degassed with argon prior to each use. Peptide substrate was dissolved in buffer to a final concentration of 1mM. The following were added: (1) 5% v/v *t*-BuSH and (2) 0.5 mg/mL titanium oxide ($TiO_2$, anatase, Thermo Fisher Scientific). The reaction vessel was sealed with parafilm, placed in a floating rack into an Bransonic 1510R-MTH Branson Ultrasonic Cleaner (Artisan Technology Group, Champaign, IL, USA) and ultrasonicated (40 kHz, 100% intensity (9 mW/$cm^3$), 37 °C). The reaction was monitored by analytical HPLC and MALDI-TOF MS. Analytical HPLC timepoints were first diluted 25-100x with Milli-Q water, degassed briefly with argon gas and injected on a Luna Omega C18 analytical column (5 µM particle size, 150 mm length, 4.6 mm diameter). MALDI timepoints were diluted by 25x in MilliQ water and co-spotted with CHCA matrix.

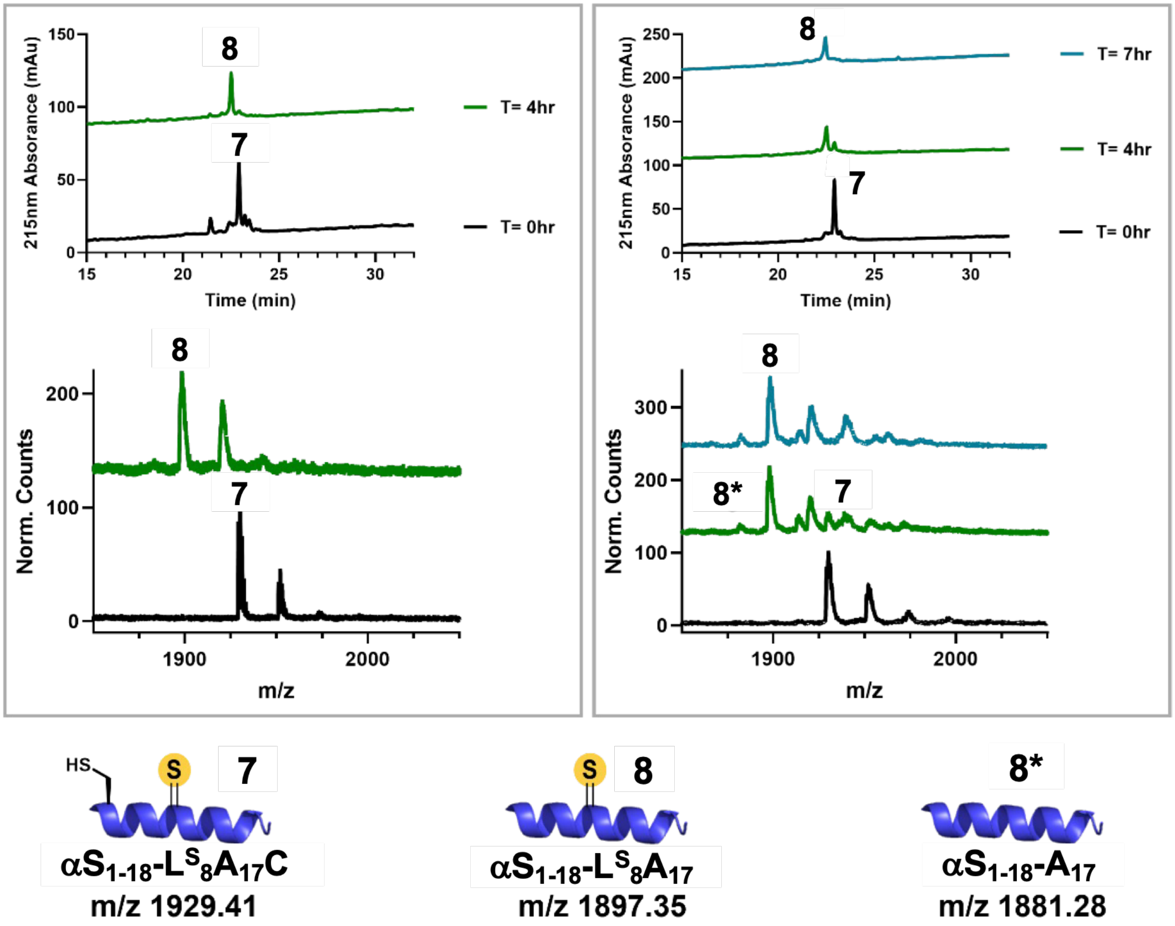


**Figure S9**. Comparison of VA-044 initiated (left) and USID (right) to desulfurize αS$_{1\text{-}18}$-L$^{S}_{8}$A$_{17}$C (**7**) to form αS$_{1\text{-}18}$-L$^{S}_{8}$A$_{17}$ (**8**). Both desulfurization reactions were performed on a 65 nmol scale. The analytical (top) and MALDI (bottom) data demonstrate complete conversion to desulfurized product in both cases. Bottom: calculated $[M+H]^{+}$ masses are shown. Thioamide desulfurization product (**8***). Calculated $[M+Na]^{+}$ are as follows: αS$_{1\text{-}18}$-L$^{S}_{8}$A$_{17}$C: 1952.4, αS$_{1\text{-}18}$-L$^{S}_{8}$A$_{17}$: 1920.4, and αS$_{1\text{-}18}$-L$_{8}$A$_{17}$: 1904.3. Calculated $[M+2Na]^{+}$ are as follows: αS$_{1\text{-}18}$-L$^{S}_{8}$A$_{17}$C: 1975.4, αS$_{1\text{-}18}$-L$^{S}_{8}$A$_{17}$: 1943.4, αS$_{1\text{-}18}$-L$_{8}$A$_{17}$: 1927.3. Observed masses are as follows: (T = 0 hr: VA-044 1929.2, 1951.2, USID: 1929.5, 1951.0, 1973.1. T = 4 hr: VA-044: 1897.5, 1919.6, USID: 1881.3, 1897.3, 1919.1, 1929.3, T = 7 hr: USID: 1881.4, 1897.4, 1919.2, 1939.2.

### Desulfurization of Ligation Site Cysteine

Additional details can be found as previously described.[1] Desulfurization buffer was prepared (6 M Gdn HCl, 0.2 M $NaHPO_4$, 0.5 M TCEP, 0.2 M thioacetamide pH 7.0; TCEP/ thioacetamide were added on the day of use and pH adjusted accordingly) and degassed with argon. The NCL product was dissolved in enough desulfurization buffer for a final concentration of 0.1 mM. The following were added: (1) 10% (v/v) *tert*-butylthiol (*t*-BuSH) and (2) 0.5 M VA-044 to achieve a final concentration of 50 mM VA-044. The reaction was sealed with parafilm, left to incubate at 37 °C, and was monitored with MALDI-TOF MS. MALDI -TOF MS timepoint aliquots were diluted by 200x with Milli-Q water with 0.1% TFA and co-spotted with CHCA matrix. After 4-5 hours, the reaction was purged with argon for 15 minutes to remove excess *t*-BuSH. The reaction was diluted with Milli-Q water and immediately purified by RP-HPLC with a C18 semi-preparative column (Luna Omega PS C18 with 5 μm particle size, 250 mm length, 10 mm diameter (Phenomenex)) with 20-40% gradient (**Table S4**, **H**).

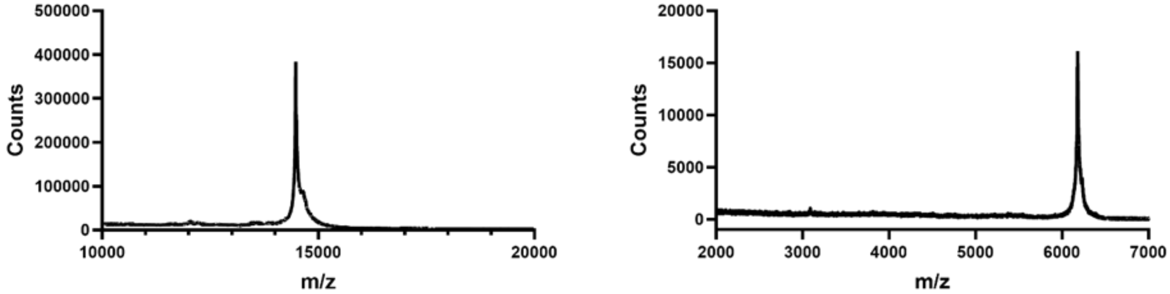


**Figure S10.** MALDI of the purified desulfurized αS-$L^{S}_{8}$ (Left) and GB1-$L^{S}_{5}K_{13}A$ (Right) constructs. **Left)** (calculated = $[M+H]^+$ 14476.22, observed = $[M+H]^+$ 14478.81). **Right)** (calculated = $[M+H]^+$ 6183.79, observed = $[M+H]^+$ 6181.63)

**Table S3.** MALDI and characterization of purified GB1 and αS protein fragments and NCL products

| Construct | Calc. $[M+H]^+$ | Obs $[M+H]^+$ | %Conversion/ % Yield |
|---|---|---|---|
| $GB1_{13\text{-}56}$-$K_{13}C$ | 4796.11 | 4790.89 | N/A |
| GB1-$K_{13}A$ | 6166.80 | 6165.84 | N/A |
| GB1-$L^{S}_{5}K_{13}C$ NCL Product | 6215.85 | 6215.73 | Pyrazole: 92%/38%<br>Azide: 95%/55% |
| GB1-$L^{S}_{5}K_{13}A$ Post-desulfurization | 6183.79 | 6181.63 | 100%/56% |
| $\alpha S_{19\text{-}140}$-$A_{19}C$ | 12629.96 | 12630.24 | N/A |
| αS-$L^{S}_{8}A_{19}C$ NCL Product | 14509.23 | 14507.89 | Pyrazole: 61%/9%<br>Azide: 65%/11% |
| αS-$L^{S}_{8}$ Post-desulfurization | 14476.23 | 14478.81 | 87%/25% |

**Table S4.** HPLC and FPLC methods for proteins, protein segments, NCL products, and post-desulfurization product purification. All HPLC methods use MilliQ water with 0.1% TFA as solvent A and acetonitrile with 0.1% TFA as solvent B. FPLC solvents are explained in the text.

| **Method** | **CV** | **% B** | **Grad.** | **CV** | **%B** |
|---|---|---|---|---|---|
| **A** | 0 CV | 15 | **B** | 0 CV | 0 |
| | 1.2 CV | 15 | | 2 CV | 0 |
| | | | | 42 CV | 40 |
| | | | | 48 CV | 100 |
| | | | | 50 CV | 0 |
| **Grad.** | **Time** | **% B** | **Grad.** | **CV** | **%B** |
| **C** | 0.00 | 5 | **D** | 0 CV | 0 |
| | 5.00 | 5 | | 2 CV | 0 |
| | 50.00 | 50 | | 28 CV | 30 |
| | 53.00 | 100 | | 32 CV | 100 |
| | 58.00 | 100 | | 34 CV | 0 |
| | 60.00 | 5 | | | |
| **Grad.** | **Time** | **% B** | **Grad.** | **Time** | **% B** |
| **E** | 0.00 | 5 | **F** | 0 CV | 0 |
| | 5.00 | 5 | | 3 CV | 0 |
| | 10.00 | 33 | | 13 CV | 20 |
| | 40.00 | 43 | | 23 CV | 35 |
| | 43.00 | 100 | | 28 | 45 |
| | 48.00 | 100 | | 34 CV | 0 |
| | 50.00 | 5 | | | |
| **Grad.** | **CV** | **%B** | **Grad.** | **Time** | **%B** |
| **G** | 0.00 | 5 | **H** | 0.00 | 5 |

| | | | | | |
|---|---|---|---|---|---|
| | 3.00 | 5 | | 10.00 | 5 |
| | 5.00 | 20 | | 15.00 | 20 |
| | 11.00 | 20 | | 55.00 | 40 |
| | 51.00 | 35 | | 58.00 | 40 |
| | 53.00 | 100 | | 65.00 | 100 |
| | 58.00 | 100 | | 70.00 | 100 |
| | 60.00 | 5 | | 75.00 | 5 |

| **Grad.** | **CV** | **%B** |
|---|---|---|
| **I** | 0.00 | 2 |
| | 10.00 | 2 |
| | 15.00 | 20 |
| | 55.00 | 40 |
| | 58.00 | 100 |
| | 65.00 | 100 |
| | 68.00 | 2 |

### Circular Dichroism

*General* The dried aliquot of GB1 protein was dissolved in 6 M Gdn HCl, 20 mM $Na_2HPO_4$ pH 7.0 (1 mL) and stirred for 2 hours at room temperature. The denaturant was diluted to 5 M Gdn HCl upon addition of 200 µL of CD buffer (20 mM $Na_2HPO_4$ pH 7.0) and allowed to stir for 30 minutes. Followed by addition of 300 µL of CD buffer (diluted to 4 M Gdn HCl) for 2 hours, 500 µL of CD buffer (diluted to 3 M Gdn HCl) for 1.5 hours, and 1 mL of CD buffer (diluted to 2 M Gdn HCl) for 1.5 hours. A final 2 mL of CD buffer was added, and the sample was dialyzed overnight into 5 L of CD buffer at 4 °C. The dialysis was switched three times and the final concentration was determined by UV-Vis ($\varepsilon_{280nm}$ = 9970 $M^{-1}$ $cm^{-1}$ (GB1-$K_{13}$A) or $\varepsilon_{274nm}$ = 19,736 $M^{-1}$ $cm^{-1}$ (GB1-$L^{S}_{5}K_{13}$A)).

*Wavelength Scan* The CD experiments were performed on a Jasco J-1500 CD spectrometer with a 1 mm path length Helma 110-QS CD cuvette. Wavelength absorbance scans were performed at 25°C, scanning from 350 to 190 nm with a continuous scanning rate of 50 nm/min (bandwidth = 1 nm and data pitch = 1 nm). The instrument was blanked with 20 mM $Na_2HPO_4$ pH 7.0 prior to sample collection. This blank was manually subtracted from the sample data. The raw signal (θ, mDeg) was converted to the mean residue ellipticity ($\theta_{MRE}$) (**Equation S1**) where l is the pathlength in cm, $n_R$ is the number of residues and c is the concentration in M.

$$\theta_{MRE} = \frac{\theta}{c * l * n_R}$$ **Equation S1**

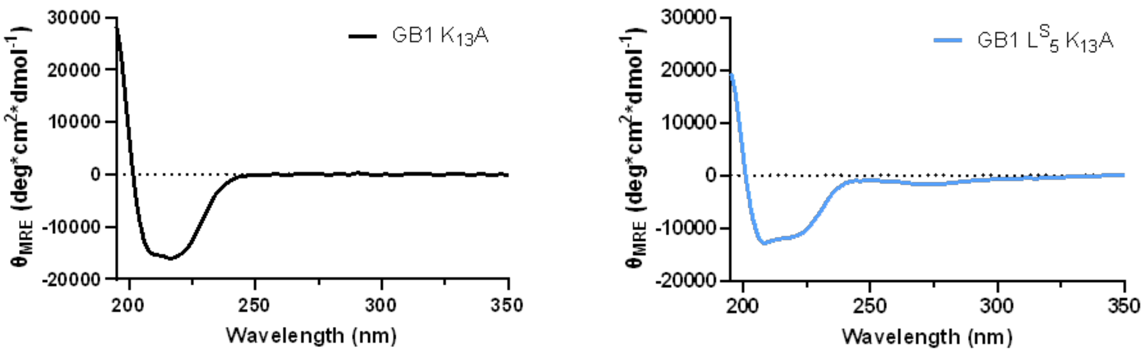


**Figure S11.** Wavelength absorbance scan of GB1-$K_{13}$A and GB1-$L^{S}_{5}K_{13}$A. The thioamide signature is apparent around 270 nm for GB1-$L^{S}_{5}K_{13}$A. GB1 $K_{13}$A appears well folded, whereas the thioamide construct has less 220 nm signature.

*Thermal Denaturation* Thermal denaturation was monitored by changes in θ (mDeg) at 220 nm from 4-6 °C to 95 °C at a rate of 0.2 °C/min with an equilibration time of 10 seconds (interval = 1 °C, DIT = 8 sec and bandwidth = 1 nm). The data was acquired in duplicate. The instrument was blanked with 20 mM $Na_2HPO_4$ pH 7.0 prior to sample collection. CD cuvettes were parafilmed to avoid evaporation at the high temperature required to thermally denature GB1. The linear folded (**Equation S2**) and unfolded (**Equation S3**) baselines were fit from 4 °C to 40 °C and 75 °C to 95 °C for GB1-$L^S{}_5K_{13}A$ and 6 °C to 60 °C and 80 °C to 95 °C for GB1-$K_{13}A$ respectively using GraphPad Prism 7.01 software (San Diego, CA, USA). The entire dataset was fit to **Equation S6** with ΔH and ΔS as adjustable parameters (T (Kelvin), R (8.314 J $mol^{-1}$ $K^{-1}$)) by using the solver function in Microsoft Excel (Redmond, WA, USA) to minimize the sum of the squared residuals. The fraction unfolded curve is generated by performing the same minimization of the sum of the squared residuals between $F_{Calc}$ and the experimental fraction unfolded ($F_{Exp}$, **Equation S7**). The resulting fits of the experimental $\theta_{MRE}$ data and fraction folded plots (1- $F_{Calc}$) generated in GraphPad Prism are displayed in **Figure S12**.

$$\theta_f = m_f T + b_f \quad \textbf{Equation S2}$$

$$\theta_u = m_u T + b_u \quad \textbf{Equation S3}$$

$$K = e^{\frac{-\Delta H + T\Delta S}{RT}} \quad \textbf{Equation S4}$$

$$F_{Calc} = \frac{K}{K+1} \quad \textbf{Equation S5}$$

$$CD_{Fit} = \theta_f(1 - F_{Calc}) + \theta_u * F_{Calc} \quad \textbf{Equation S6}$$

$$F_{Exp} = \frac{\theta_{MRE} - \theta_f}{\theta_u - \theta_f} \quad \textbf{Equation S7}$$

The other values derived from the thermal denaturation experiments are displayed in **Table S5**. $T_m$, $\Delta G_{25}$ and $\Delta\Delta G_U$ are calculated using **Equation S8**, **Equation S9** (T = 298.15 K), and **Equation S10** respectively.

$$T_m = \frac{\Delta H}{\Delta S}$$ **Equation S8**

$$\Delta G = \Delta H - T\Delta S$$ **Equation S9**

$$\Delta\Delta G_U = \frac{\Delta T_m * \Delta H_{Oxo}}{T_{m,Oxo}}$$ **Equation S10**

**Table S5.** Thermodynamic parameters calculated from the two-state fits of the thermal denaturation data.

| Construct | $T_m$ (°C) | ΔH (kJ mol$^{-1}$) | ΔS (kJ mol$^{-1}$ K$^{-1}$) | $\Delta G_{25}$ (kcal mol$^{-1}$) | $\Delta\Delta G_U$ (kcal mol$^{-1}$) |
|---|---|---|---|---|---|
| GB1-$K_{13}$A | 68.2 ± 0.34 | 311 ± 6.7 | 0.91 ± 0.02 | 9.41 ± 0.14 | --- |
| GB1-$L^S{}_5K_{13}$A | 60.7 ± 0.03 | 198 ± 2.1 | 0.59 ± 0.01 | 5.06 ± 0.05 | -1.6 |

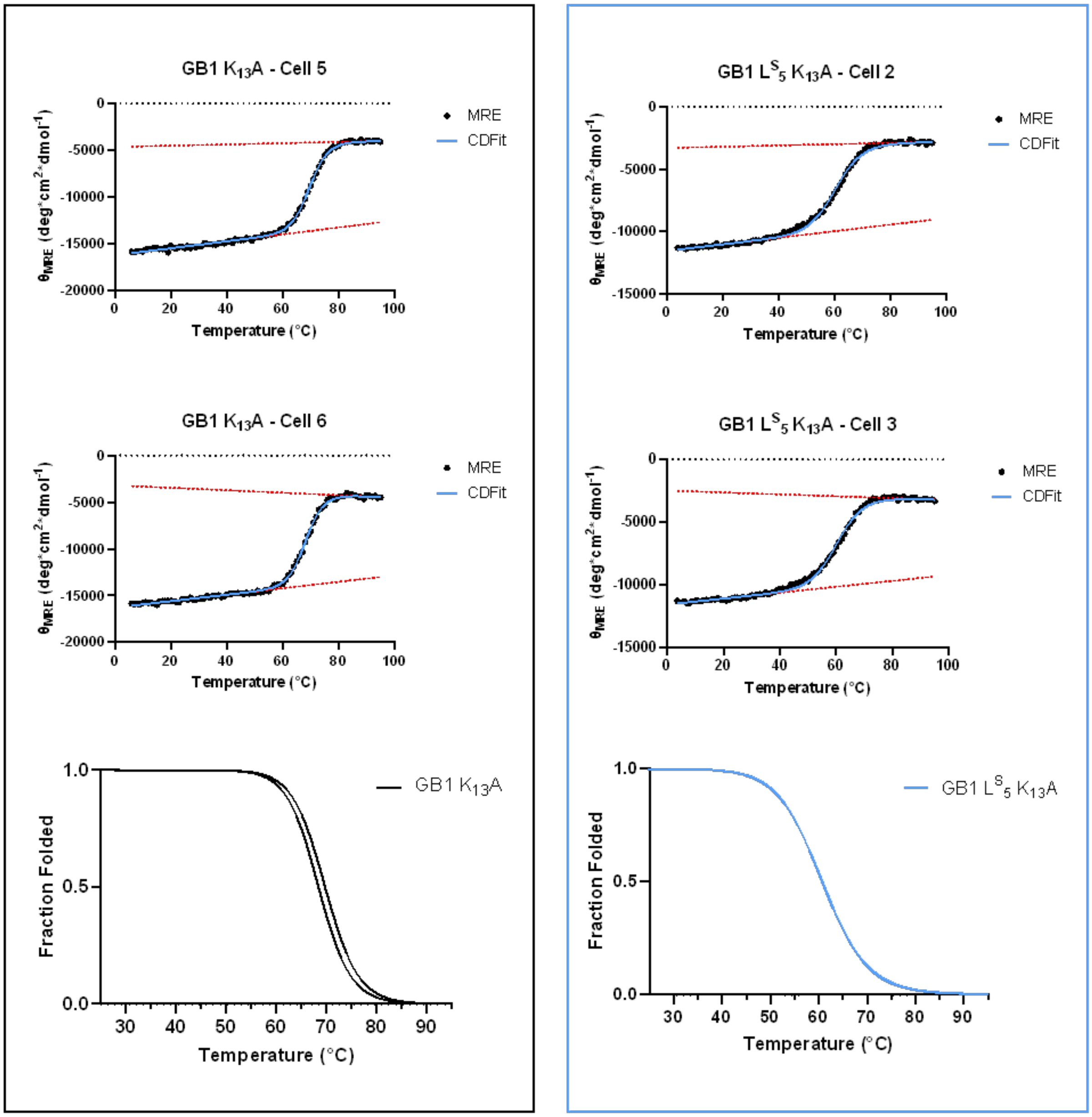


**Figure S12.** Raw (θMRE) values from thermal denaturation experiment fitted to a two-state unfolding (CDFit) for GB1-$K_{13}$A and GB1-$L^S_5K_{13}$A. The duplicate data is shown for GB1-K13A (left) and GB1-$L^S_5$ K13A (right). The unfolded and folded baselines are in red. The fraction folded plots (1-FCalc) for both replicates are displayed.

**Crystallization and Structure Determination of GB1-$K_{13}$A**

Crystals of GB1-K13A were grown by the sitting drop vapor diffusion method at 4 °C. A 100 nL drop of protein solution [10 mg/mL GB1-$K_{13}$A in 50 mM phosphate buffer (pH 5.5)] was added to 100 nL of precipitant solution [2.0 M ammonium citrate tribasic (pH 7.0), 0.1 M BIS-TRIS propane (pH 7.0)] and equilibrated against a 50 µL reservoir of the precipitant solution. Crystals of GB1-K13A were briefly immersed in cryoprotectant solution (mother liquor supplemented with 20% ethylene glycol) before being flash-cooled in liquid nitrogen.

X-ray diffraction data were collected on beamline 9-2 at the Stanford Synchrotron Radiation Laboratory (SSRL) at Stanford University. Diffraction data were indexed and integrated using iMosflm[7] and scaled using Aimless[8] in the CCP4 program suite.[9] The crystal structure of GB1-K13A was determined by molecular replacement using Phaser[10] and the X-ray crystal structure of WT GB1 (PDB 2QMT)[11] as the search model for rotation and translation function calculations. Iterative cycles of refinement and manual model building were performed using Phenix[12] and WinCoot,[13] respectively. Refinement proceeded until $R_{free}$ converged at its lower limit. The quality of the final model was assessed using MolProbity.[14] Data collection and structure refinement statistics are listed in **Table S6**.

The crystal structure of GB1-K13A was determined at 1.37 Å resolution (**Figure S13**) with excellent refinement statistics (**Table S6**). Electron density for the A13 side chain was well-defined. The K13A substitution did not perturb the main chain conformation and the root-mean-square deviation (RMSD) of 56 C$\alpha$ atoms between WT GB1 (PDB 2QMT) and GB1-K13A was calculated as 0.65 Å by Superpose (**Figure S13**).[15] Atomic coordinates and structure factor amplitudes have been deposited in the Protein Data Bank (www.rcsb.org) with accession code 12LO.

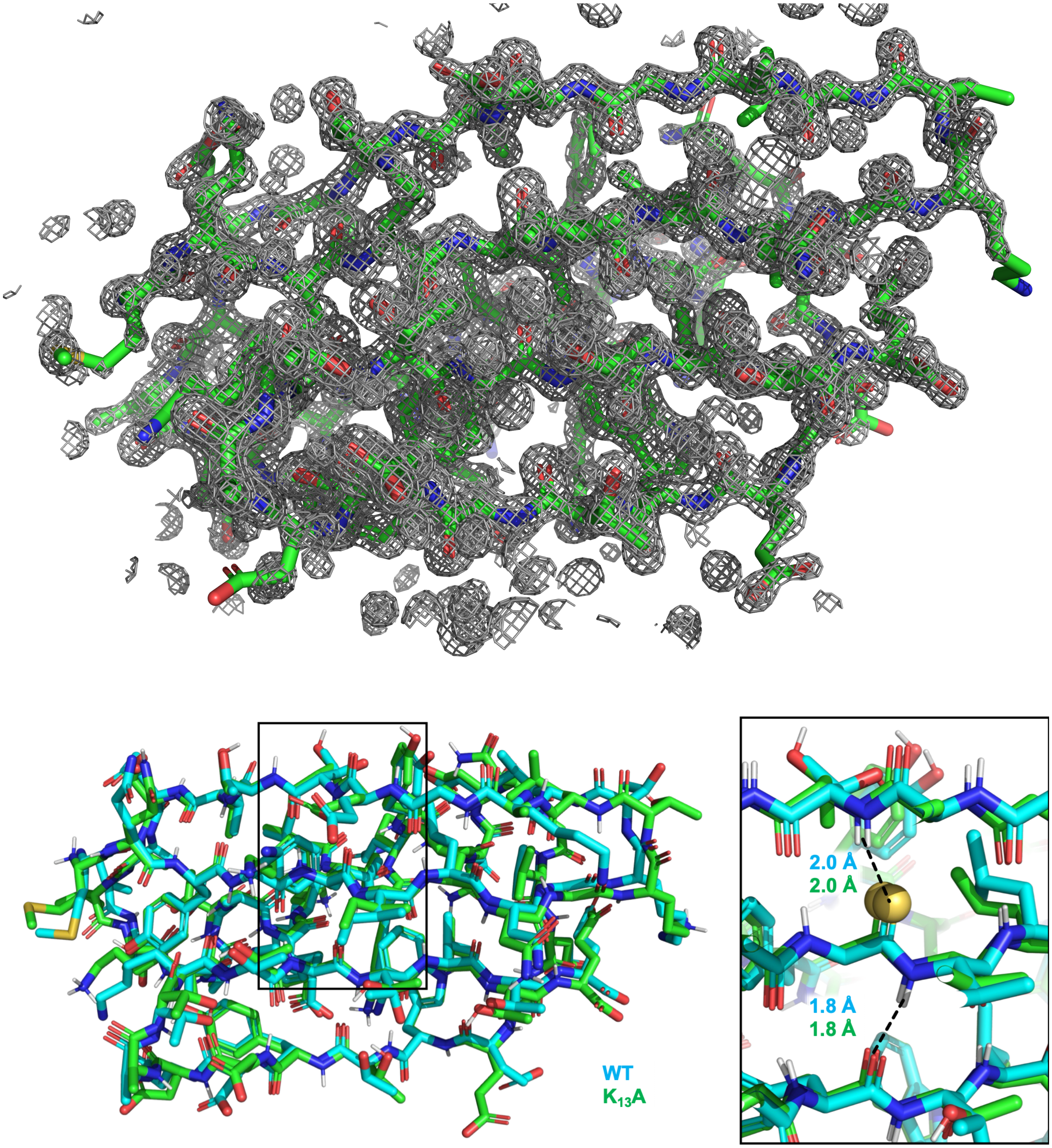


**Figure S13.** GB1-$K_{13}$A X-ray Structure. Top: 2f0-fc map of GB1-$K_{13}$A. Bottom: Structures of WT GB1 (teal) and GB1-$K_{13}$A (green). Backbone C$\alpha$ alignment of WT GB1 structure (PDB 2QMT)[11] and GB1-$K_{13}$A structures shows that the GB1 fold is preserved in GB1-$K_{13}$A with the same positioning and H-bonding distances for $L_5$.

**Table S6.** GB1-$K_{13}$A Data Collection and Refinement Statistics

| **Unit Cell** | |
|---|---|
| **Space group** | $C222_1$ |
| ***a, b, c*** **(Å)** | 42.28, 80.01, 31.74 |
| **α, β, γ (deg)** | 90, 90, 90 |
| **Data Collection** | |
| **Resolution (Å)[a]** | 40.0 – 1.37 (1.39 – 1.37) |
| **Total no. of reflections[a]** | 68728 (3413) |
| **No. of unique reflections[a]** | 11615 (553) |
| **Multiplicity[a]** | 5.9 (6.2) |
| **Completeness (%)[a]** | 98.9 (97.2) |
| **$R_{merge}$[a]** | 0.068 (0.402) |
| **$R_{pim}$[a]** | 0.030 (0.174) |
| **$CC_{1/2}$[a]** | 0.998 (0.897) |
| **I/σ(I)[a]** | 13.3 (4.0) |
| **Refinement** | |
| **No. of reflections used in refinement/test set** | 11606/581 |
| **$R_{work}$** | 0.1721 |
| **$R_{free}$** | 0.2340 |
| **Rmsd from ideal geometry** | |
| **Bonds (Å)** | 0.005 |
| **Angles (deg)** | 0.8 |
| **Ramachandran plot (%)** | |
| **Favored** | 98.15 |
| **Allowed** | 1.85 |
| **Outliers** | 0 |
| **MolProbity score** | 0.832 |

[a] Values in parentheses refer to the highest-resolution shell of the data.

### $^{1}H$-$^{1}H$ TOCSY NMR

NMR data were collected on a Bruker AVANCE NEO 600 MHz spectrometer. $^{1}H$-$^{1}H$ TOCSY was performed by collecting 4096 points in *f*2, 512 points in *f*1 with 32 scans at 298 K. The water $^{1}H$ signal (ppm) was determined for each sample and inputted as the Transmitted frequency offset (OP1). The spectra were processed with MestReNova 14.1.0 (Santiago de Compostela, Spain). Apodization of Sine Square 90˚ was used for both *f*2 and *f*1, and zero-fill was 2x the size of the FID. A baseline correction of a Berstein polynomial fit of order 3 was used.

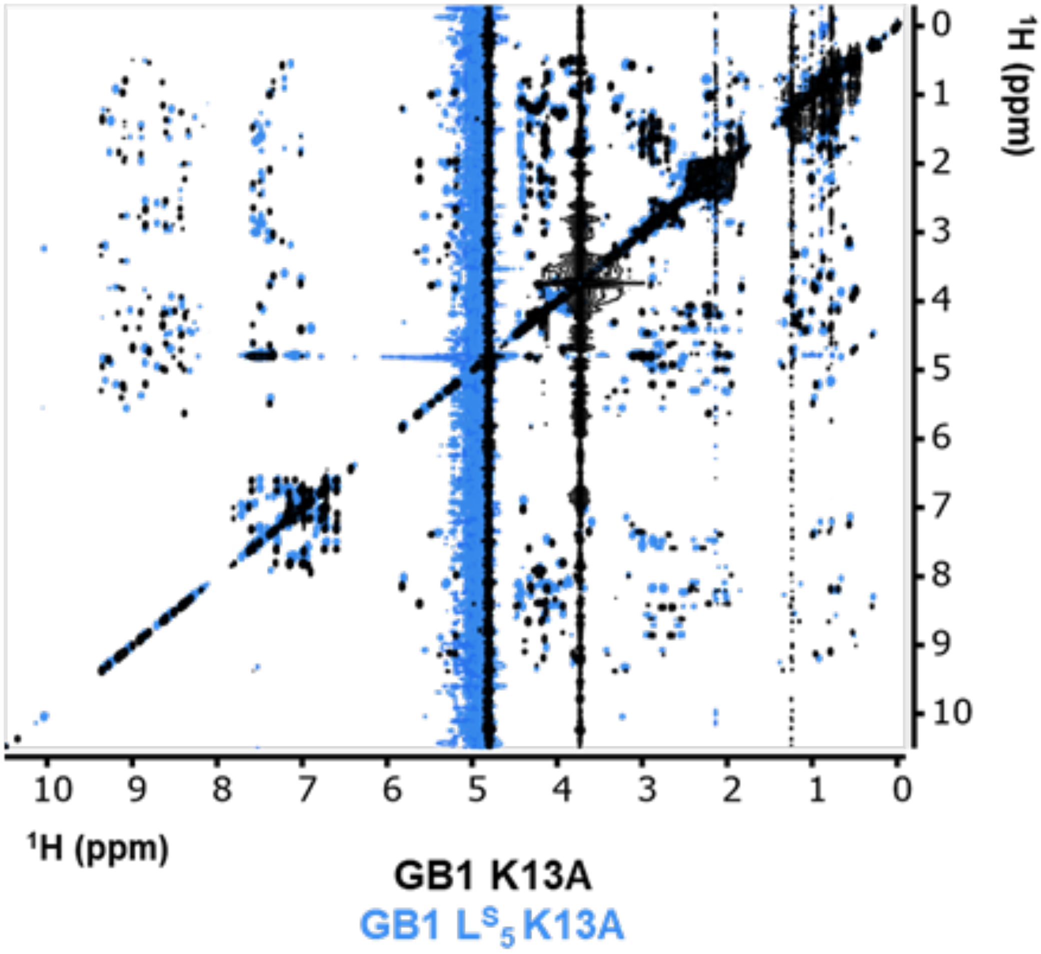


**Figure S14.** Overlay of $^{1}H$-$^{1}H$ TOCSY spectra for GB1-$K_{13}A$ and GB1-$L^{S}{}_{5}K_{13}A$.

### αS Aggregation Kinetics

To form pre-formed fibrils (PFFs), a 100 μM stock of unmodified αS in 1x phosphate buffered saline (PBS) was shaken at 37°C for 3-7 days. Dried aliquots of αS-$L^S$8 were dissolved in PBS to 100 μM. A 1 mM stock of thioflavin-T (ThT) in 1x PBS was prepared and stored at -20 °C until use. For each percentage of thioamide αS monomer, stock solutions of thioamide-containing αS monomer, unmodified αS monomer and ThT were prepared (0%: 40 μM unmodified αS monomer and 10 μM ThT, 25%: 30 μM unmodified αS monomer, 10 μM thioamide-containing αS monomer and 10 μM ThT, 50%: 20 μM unmodified αS monomer, 20 μM thioamide-containing αS monomer and 10 μM ThT, 100%: 40 μM thioamide-containing αS monomer and 10μM ThT). 50 μL of each stock solution was pipetted into a 96-well plate (Greiner Bio-One, microplate, 96 well, PS, half area, black) in triplicate. The plate was sealed and shaken in a 37 °C at 1400 rpm for 15 minutes. To make unmodified αS seeds, the PFF stock solution was diluted to 4 μM in 1x PBS. The diluted PFF stock solution was then placed in an ice water bath and sonicated at amplitude 50 for 2 minutes (1s on, 1s off). 50 μL unmodified αS seeds were then added to the well plate to give final concentrations of 2 μM seeds, 20 μM αS monomer and 5 μM ThT. Well plate was sealed tightly and shaken at 37 °C at 1400 rpm. ThT fluorescence emission was monitored using a Spark Mulitmode Microplate Reader (Tecan Trading AG; Switzerland) at 485 nm (excitation at 440nm) every few hours until aggregation was complete.

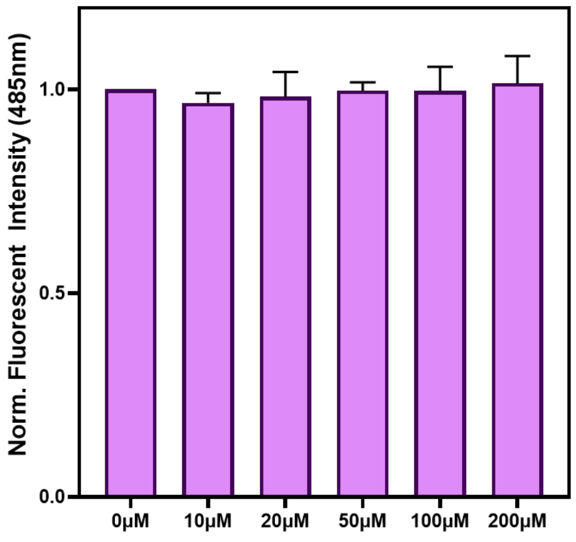


**Figure S15.** Normalized fluorescent emission of ThT in the presence of thioacetamide. 100μL stocks containing 20μM αS PFFs 10μM ThT in 1x PBS were made for each thioacetamide concentration. Each concentration of thioacetamide was examined in triplicate. ThT fluorescence emission was monitored using a Spark Mulitmode Microplate Reader (Tecan Trading AG; Switzerland) at 485nm (excitation at 440nm).

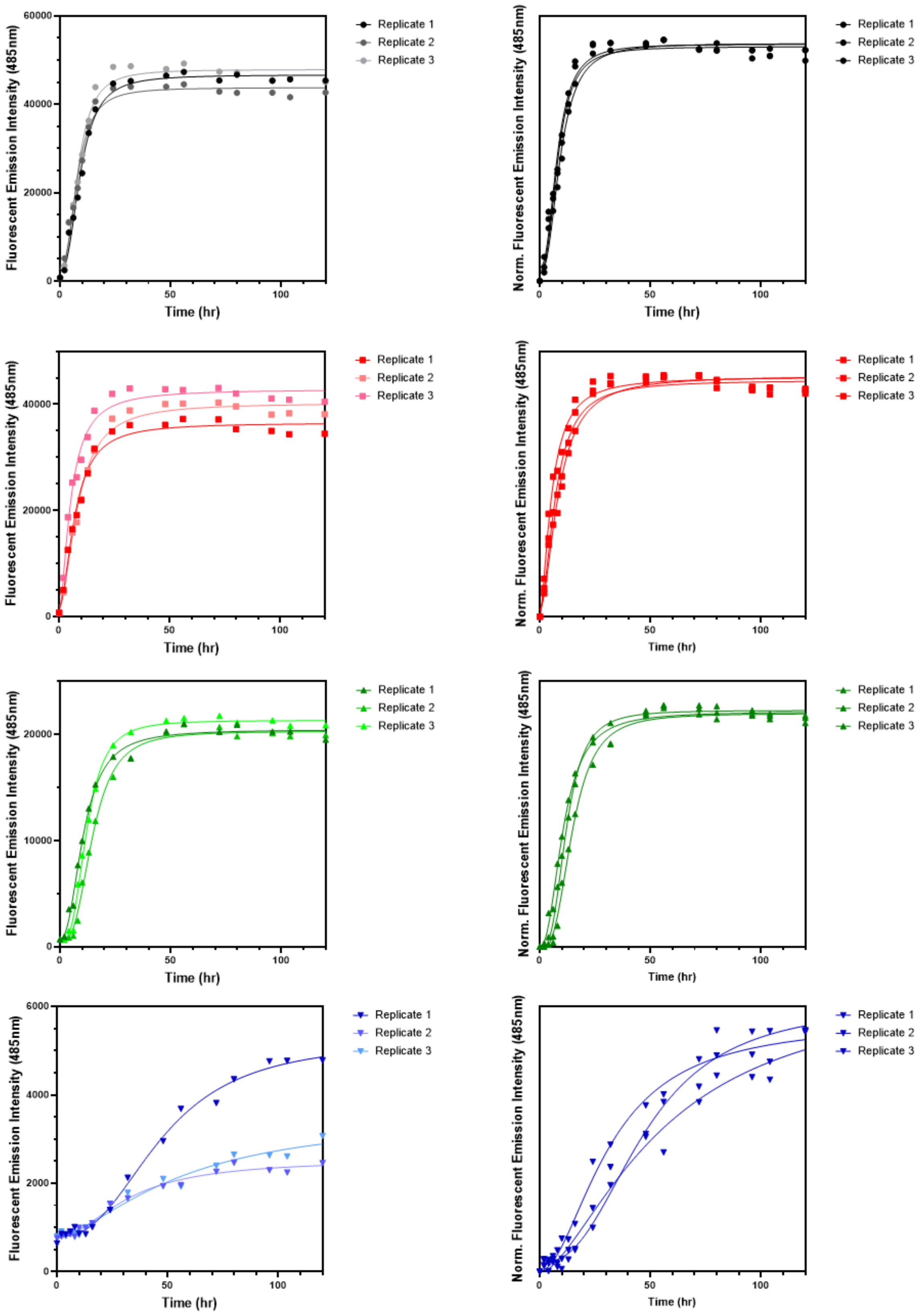


**Figure S16.** ThT aggregation kinetics assays with different percentages of αS-L$^{S}{}_{8}$. Unnormalized (Left) and normalized (Right) aggregation kinetics of 0% (black), 25% (red), 50% (green) and 100% (blue) αS-L$^{S}{}_{8}$.

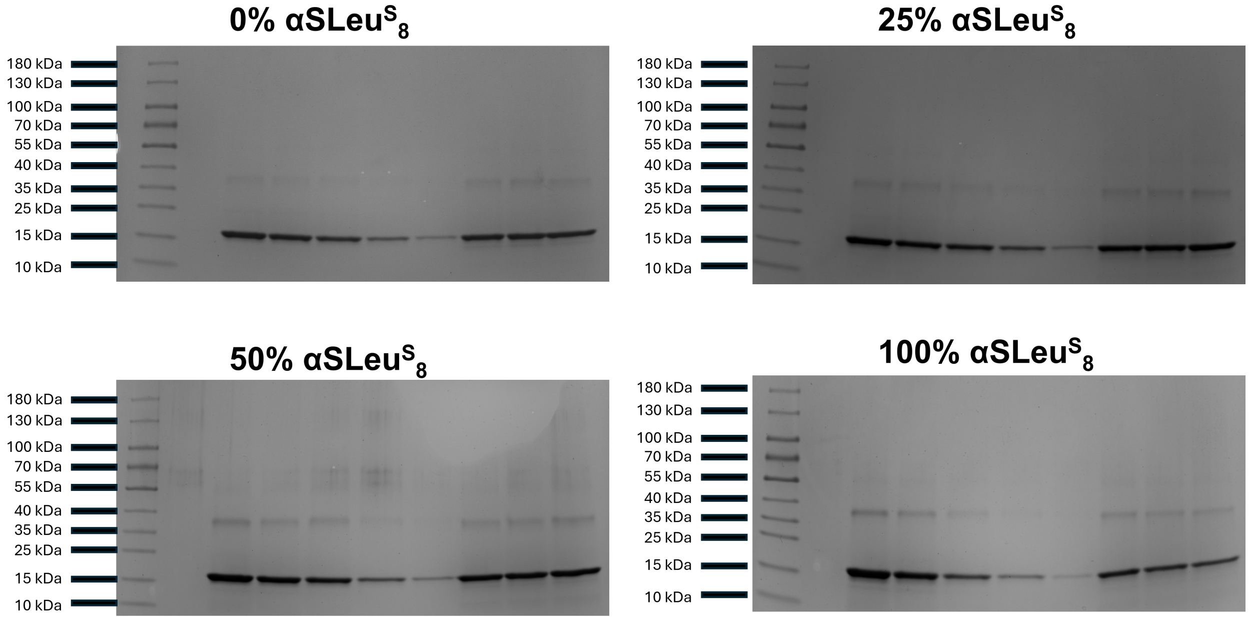


**Figure S17.** 12-18% Tris-glycine acrylamide SDS-PAGE gels of monomer incorporation assay of αS into fibrils during aggregation kinetics assay. Gel was stained with Coomassie sensitive stain (50 g aluminum sulfate, 100 mL ethanol, 200 mg Coomassie Brilliant Blue G-250, 23.5 mL O-phosphoric acid and 800+ mL of Milli-Q water for a final volume of 1000mL). The left lane on gels represents ladder standard (PageRuler™ Prestained Protein Ladder, 10 to 180 kDa (Thermo Fisher)). Next five lanes following represent monomeric αS protein standards (20 μM, 15 μM, 10 μM, 5 μM and 2.5 μM).

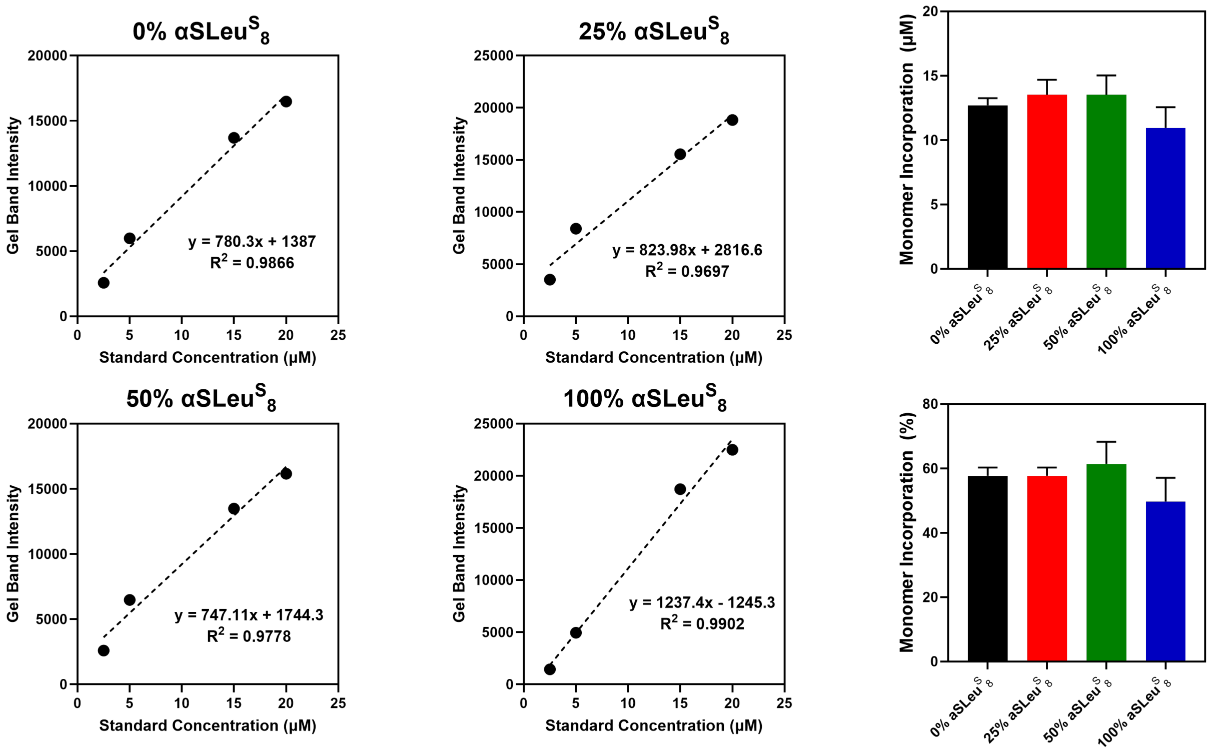


**Figure S18.** Quantification of monomer incorporation during ThT aggregation kinetics assay αS fibrilization. Quantifying gel band densities was performed using ImageJ. All monomer incorporation differences were determined non-significant compared to 0% αS-L$^S_8$ ($p > 0.5$) by unpaired t-test.
